# Changes in peripheral blood leukocyte composition precede development of heart-reactive autoantibodies in patients hospitalised for acute heart failure

**DOI:** 10.1101/2025.11.01.685879

**Authors:** Boshra Afshar, Maximilian Bauser, Dora Pelin, Sahel Shemshadi, Claudia Hollmann, Hanna S. Hepp, Wafaa Al Hassan, Mairin Heil, Dennis Göpfert, Lena Schmidbauer, Elisa Kaiser, Nele Gellermann, Jannika Pätkau, Fabian Kerwagen, Peter Heuschmann, Georg Gasteiger, Wolfgang Kastenmüller, Gustavo Ramos, Thomas Kerkau, Ulrich Hofmann, Stefan Frantz, Stefan Störk, Caroline Morbach, Niklas Beyersdorf

**Author notes:** Correspondence: Niklas Beyersdorf, MD University of Würzburg, Institute for Virology and Immunobiology Versbacher Str. 7, 97078 Würzburg Germany.

## Abstract

In a retrospective pilot study, we showed that the induction of heart-reactive autoantibodies (HRA) in the wake of acutely decompensated heart failure predicts worse outcomes. To gain deeper insights into the immunological mechanisms causing induction of HRA after heart failure decompensation we initiated the prospective ‘Acute Heart Failure-Immunomonitoring Cohort Study’ (AHF-ImmunoCS). For this study, 380 patients were enrolled and will be followed up, including serial collection of biomaterials, for a period of 18 months after the index hospitalisation for AHF. Analysis of AHF-ImmunoCS samples obtained at baseline and at 6-month follow-up from 110 patients showed *de novo* induction of HRA - as detected by indirect immunofluorescence (IFT) - in 21% of patients (previously published percentage: 32%). The IFT results did not reflect induction of broad anti-heart autoimmunity as autoantibodies against other cardiac antigens like Troponin I3 or Myosin Light Chain 7 were not induced in parallel. To understand what drives HRA induction in these patients we longitudinally immunophenotyped peripheral blood leukocytes at baseline, 6-week and 6-month follow-up by high-resolution spectral flow cytometry. Among lymphocytes, induction of HRA in the wake of acute decompensation of heart failure was associated with a higher proportion of CD4^+^ T cells among lymphocytes, more CD45RA^+^ CCR7^+^ naive conventional, i.e. non-regulatory, and more CXCR3^+^ CCR4^-^ Th1 cells among CD4^+^ T cells at baseline. Among myeloid cells, there were no differences at baseline between patients going on to develop HRA and those that did not. However, patients developing HRA had higher proportions of eosinophils (six-month follow-up) and lower proportions of Arginase^+^ HLA-DR^-^ polymorphnuclear myeloid-derived suppressor cells among myeloid cells (six-week and six-month follow-up). Our data, thus, implicate that alterations in the composition of both the lymphoid and the myeloid compartments might drive HRA induction which impacts disease progression and prognosis in AHF.

## Introduction

Acute heart failure (AHF) is characterised by heart failure symptoms that require medical intervention and usually leads to patient hospitalisation. AHF can be due to new onset (de novo heart failure (HF)) or worsening of preexisting HF and is the leading cause of unexpected hospital admissions in patients over 65 years old (1). Approximately 10% of patients with AHF do not survive hospitalisation, and nearly one-third dies within a year of an AHF episode. Despite advancements in chronic HF management, no novel therapeutic approaches for AHF have been developed in the past two decades (1, 2). Consequently, AHF is a significant public health concern, a substantial financial burden, and a challenge for modern cardiovascular research (3).

Heart failure is characterized by the heart’s inability to generate sufficient cardiac output to meet the metabolic requirements of body for oxygen and nutrients (4). Heart failure presents in distinct phenotypes based on the ejection fraction: preserved (HFpEF, >50%), mid-range (HFmrEF, 40– 49%), and reduced (HFrEF, <40%). HFpEF is associated with metabolic and inflammatory comorbidities, structural remodelling, and impaired ventricular relaxation, whereas HFrEF results from cardiomyocyte loss leading to systolic dysfunction. HFmrEF represents an intermediate and often transitional state between these two forms (5).

Growing evidence suggests that autoimmune mechanisms play a pivotal role in the initiation and progression of HF, not only in the context of systemic autoimmune diseases but also in non-ischaemic cardiomyopathies exhibiting autoimmune characteristics (6–8). Autoimmune mechanisms in HF can manifest through the production of heart-specific autoantibodies, activation of autoreactive T cells, and persistent low-grade inflammation. These immune responses may be initiated by cardiac injury induced by endogenous and exogenous factors (including viral infections), leading to the exposure of previously hidden cardiac antigens and the subsequent breakdown of self-tolerance. Mechanisms such as molecular mimicry and antigenic cross-reactivity are thought to contribute significantly to the development of autoimmune responses, particularly in the context of cardiotropic viral infections (9). Additionally, systemic autoimmune diseases like Systemic Lupus Erythematosus, Rheumatoid Arthritis and Primary Sjogren’s Syndrome are associated with increased cardiovascular risk, further supporting a connection between autoimmunity and cardiac pathology (6).

Numerous studies have demonstrated that circulating autoantibodies targeting a range of cardiac self-antigens are closely associated with the development and progression of HF. Among these, autoantibodies directed against the β₁-adrenergic receptor (β₁-AR), myosin heavy chain (MyHC), and troponin have been most extensively investigated. Autoantibodies against the β₁-AR can act as receptor agonists, leading to chronic overstimulation, maladaptive intracellular signalling, cardiomyocyte apoptosis, and adverse cardiac remodelling, thereby contributing to both acute and chronic cardiac dysfunction (9–11).

In a retrospective study, we observed that the induction of heart-reactive autoantibodies (HRA) - as determined by indirect immunofluorescence on heart muscle slides (IFT) - following an episode of AHF was associated with worse clinical outcome (12). To further elucidate the immunological mechanisms underlying HRA generation after HF decompensation, we initiated the prospective Acute Heart Failure–Immunomonitoring Cohort Study (AHF-ImmunoCS) (13). This study follows 380 patients for 18-months after the index hospitalisation for AHF with serial collection of biomaterials to enable detailed longitudinal analyses of immunological parameters.

To better understand the degree of and the mechanisms behind HRA induction, we now combined detection of HRA by indirect immunofluorescence on heart tissue slices with detection of HRA with specificity for defined (cardiac) antigens as well as in-depth cellular immunophenotyping.

## Methods and Materials

### Patients and Controls

The Acute Heart Failure Immunomonitoring Cohort Study (AHF-ImmunoCS) is a prospective, monocentric cohort study conducted within the framework of the Collaborative Research Centre CRC 1525. The study complies with Good Clinical Practice guidelines and has received approval from the institutional ethics committee (#112/21). Prior to participation, all patients provide written informed consent. Enrolment occurred consecutively during hospital admission for AHF, with comprehensive phenotyping performed during the initial hospitalisation. Follow-up visits were scheduled for six weeks, six, twelve- and 18-months post admission to the index admission. The rationale and detailed study design have been described previously (13).

The reference (control) group consisted of samples of sex-and aged-matched participants of the STAAB study (Characteristics and Course of Heart Failure Stages A/B and Determinants of Progression), a prospective, population-based cohort conducted in Würzburg, Germany.This study included a comprehensively phenotyped, representative sample of 5,000 individuals aged 30 to 79 years, all free of symptomatic heart failure at baseline. Detailed study design and baseline characteristics have been described previously (14–16).

### Detection of HRA by immunofluorescence test

To detect antibodies against myocardial antigens, i.e. HRA, commercial kits were used (INOVA or Bio-Diagnostics). In brief, patient samples and samples from healthy controls were diluted either 1:20 (INOVA) or 1:8 (Bio-Diagnostics) in phosphate-buffered saline (PBS) in wells of a 96-well V-Bottom plate. Diluted sera were then incubated on the test fields for 30 min in a humidified chamber at room temperature. The slides were then gently rinsed with PBS and placed in a PBS-filled washing cuvette for 5 min. This step was repeated using fresh PBS. After drying the slides without destroying the tissue, 10 µl of FITC-conjugated anti-human IgG antibody was applied and incubated for 30 min, again in a humidified chamber at room temperature. After another round of washing, we applied mounting medium and coverslips to the slides. The samples were then independently assessed by three different observers at 400x magnification (Leica, Wetzlar, Germany) and the sample was then assigned the median of these assessments for statistical analyses.

### Bead assays for detection of HRA against defined antigens

Dynabeads™ M-270 Epoxy beads (Thermo Fisher Scientific, Waltham, USA) were coated with different sarcomeric proteins. The investigated antigens included the β1-EC_II_-AR peptide (custom-synthesised), a scrambled control peptide (custom-synthesised), Troponin I3 (TPI3) (Recombinant Human Cardiac Troponin I,), Myosin Light Chain 7 (MYL7) (Recombinant Human MYL7), Tropomyosin 1 (TPM1) (Recombinant Human Tropomyosin-1), Myosin Heavy Chain 6 (MYH6) (Recombinant Myosin Heavy Chain 6, Cardiac Muscle, Alpha), and Myosin Heavy Chain 7 (MYH7) (Fig. 1, Suppl. Fig. S3, Suppl. Table 4).

**Fig. 1.**
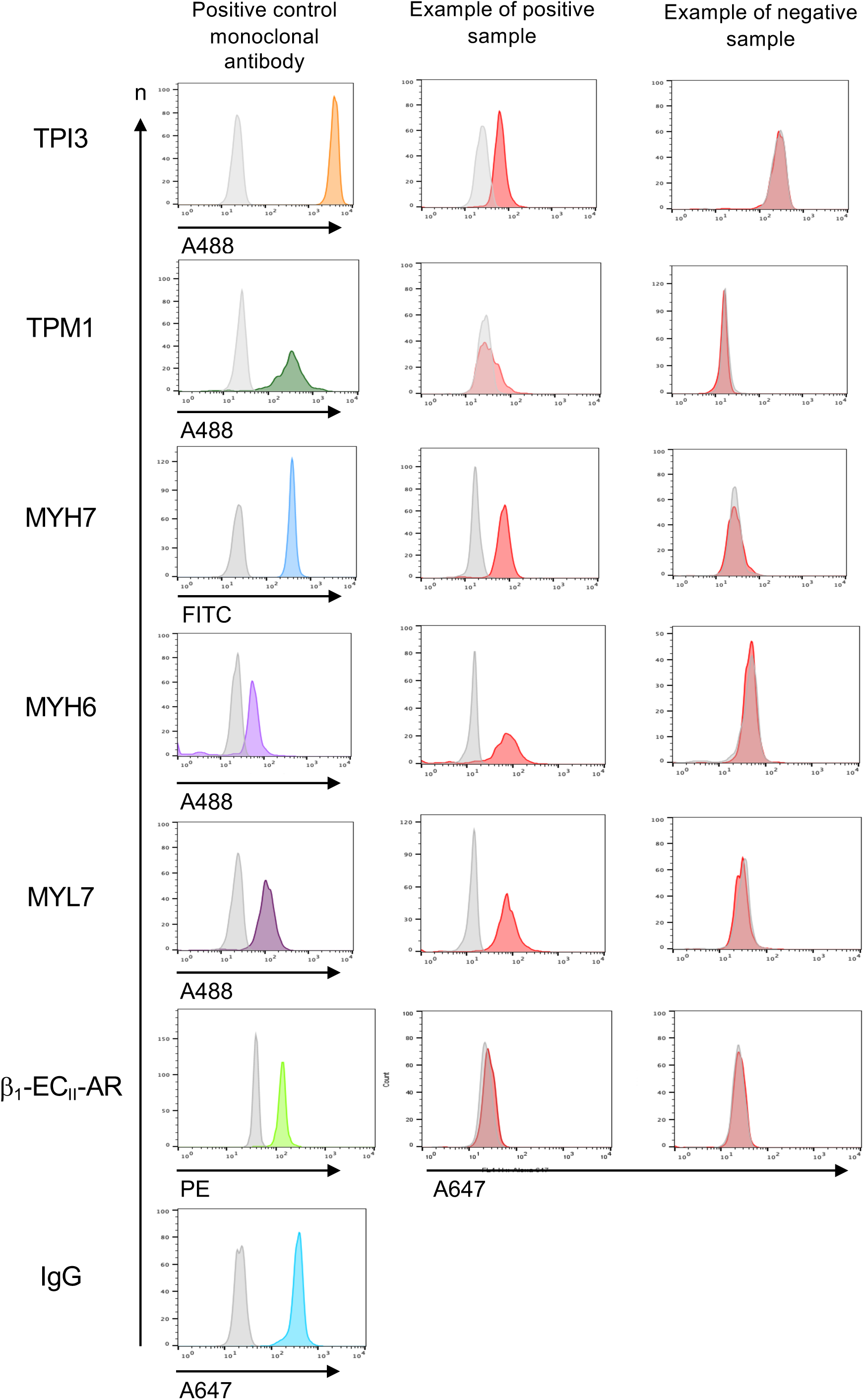
Detection of antigen-specific antibodies using protein-coated epoxy beads. Epoxy Dynabeads^TM^ were coated with target proteins or left uncoated, then incubated with human serum samples. Antibody binding was assessed by flow cytometry, with increased fluorescence intensity indicating the presence of antigen-specific antibodies. Overlay histograms show fluorescence profiles of coated beads with corresponding protein and uncoated beads, for positive control and a representative positive serum sample and negative control. A clear shift in fluorescence intensity in the coated beads relative to uncoated beads indicates antibody binding: TPI3, Troponin I3, TPM1, Tropomyosin 1, MYH7, Myosin Heavy Chain 7, MYH6, Myosin Heavy Chain 6, MYL7, Myosin Light Chain 7, β_1_-EC_II_-AR, second extracellular loop of the β1 adrenergic receptor, IgG. A488, Alexa Fluor^TM^ 488, A647, Alex Fluor^TM^ 647, FITC, Fluorescein Isothiocyanate, PE, Phycoerythrin.

A volume of 200 µL of Buffer A (0.1 M Na-phosphate buffer, pH 8) was added to each tube containing 2 x 10 ^7^ particles, mixed thoroughly and placed on a magnetic separator. The supernatant was discarded, and the washing step was repeated three times. After each addition of Buffer A, the mixture was vortexed to ensure proper suspension. A total of 70 µL of Buffer A (0.1 M Na-phosphate buffer, pH 8) and 70 µL of Buffer B (3 M ammonium sulfate, pH 7.4) were added to each tube and mixed. Subsequently, 70 µL of protein or peptide solution in PBS or 70 μl of PBS only (negative control beads) were added after thorough vortexing. The tubes were incubated on a rotator at 4°C for 72 hours, with vortexing performed once daily during incubation. On the third day, the tubes were vortexed, placed on a magnetic separator, and the supernatant was discarded. A washing step was performed by adding 200 µL of 0.1% BSA in PBS, which was repeated four times. Finally, the beads were resuspended in 200 µL of FACS buffer (phosphate-buffered saline (PBS)/ 0.1% Bovine Serum Albumin (BSA)/ 0.02% NaN_3_). On the day of analysis, beads were diluted with dilution factor of 1/100 and added to wells of a 96-well plate and washed three times with FACS buffer (centrifugation at 515 x g for 3 min; total volume per well: 200 μl) for removing unbound antigen. After discarding the buffer, 25 μl of 10% BSA in PBS was added to each well for blocking followed by incubation for 15 minutes at 4°C. Thereafter, 25 µl of serum, diluted 1:100 in FACS buffer, were added and incubated for 15 minutes at 4°C. After this step, a washing step was performed to remove any unbound serum components. Bound patient-derived antibodies were detected by addition of a secondary antibody (Alexa Fluor 647-conjugated AffiniPure Goat Anti Human IgG (H+L) Jackson ImmunoResearch Laboratories Inc, Cambridgeshire, UK). To ensure that the beads were (still) efficiently coated with antigen, a positive control consisting of an antibody specific to the coated protein was included in each run (Suppl. Table 5, 6). The reactivity of the solution containing the diluted secondary antibody was tested for each run in parallel using IgG-coated beads as the substrate. From each samples duplicates were reacted in parallel wells and measured in duplicates using a FACS Calibur flow cytometer (BD Biosciences,USA) and the data were analysed with FlowJo software (BD Biosciences,USA).

To quantitate, the fluorescence signals we calculated the ratios of the median fluorescence intensity (MFI) of coated beads to that of negative control beads using MFI means of the duplicate measurements. One was then subtracted from this ratio and the results multiplied by 100 to obtain arbitrary units of binding. Values smaller than one were set to one.

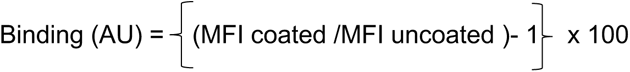

### FACS staining of patients’ whole-blood samples

10% DMSO was added to whole-blood samples prior to freezing and storage in liquid nitrogen by the interdisciplinary bank of biological materials and data Würzburg (ibdw). Samples from AHF-ImmunoCS patients were requested from the ibdw for this study. Samples were thawed in a 37° C water bath by adding PBS dropwise up to about tenfold sample volume, followed by a centrifugation step (515 x g, 10°C for 5 min). After discarding the supernatant, the pellet was resuspended in 200 µl of PBS containing a live-dead dye (Thermofisher Scientific, USA, final concentration 1:200). After 15 min of incubation at room temperature, 100 µl of the cell suspension were each aliquoted into two tubes for subsequent staining with the lymphoid antibody panel (Suppl. Table 2) or the myeloid antibody panel (Suppl. Table 3). Samples were washed by first adding 500 µl of FACS buffer followed by centrifugation (515 x g) for 5 min at 10° C.

Pellets were resuspended in 100 µl of a master mix of antibodies against surface markers (dilution in brilliant stain buffer (BD Horizon), (Suppl. Tables 2, 3) followed by a 15 min incubation at 4°C. After further washing steps with PBS to remove excess extracellular protein, pellets were resuspended in 100 µl of Fix/Perm solution (Thermofisher Scientific,USA) and incubated for 30 min at room temperature. After washing with Perm solution (Thermofisher Scientific,USA), the pellet was resuspended in 100 µl of a master mix containing antibodies with specificity for intracellular antigens, diluted in perm buffer, were added and incubated for 45 min at room temperature. After washing, the samples were resuspended in FACS buffer for measuring on the Cytek Aurora Flow cytometer (Cytek Biosciences). Comprehensive information on all antibodies used in this study is provided in Suppl. Table 2, 3. Antibody staining panels were designed and validated using a Cytek Aurora spectral flow cytometer equipped with five lasers (355, 405, 488, 561, and 640 nm; Cytek Biosciences).

### FACS data analysis

Unmixed flow cytometry data were imported into the OMIQ software platform (app.omiq.ai) for analysis. Dimensionality reduction and clustering analyses were performed using UMAP and FlowSOM algorithms within the OMIQ environment. Lymphocytes, i.e. CD8⁺ T cells, CD4⁺ T cells, B cells and NK cells and myeloid were analysed separately (Suppl. Fig. S5) UMAP analysis for lymphocytes and myeloid cells was performed with nearest neighbours of 15, a minimum distance of 0.4, the number of components set to two and using Euclidean metrics. The resulting UMAP embedding served as the basis for downstream FlowSOM clustering within the OMIQ platform. FlowSOM clustering was performed using the following parameters: 0 xdim = 12, ydim = 12, rlen = 10, random seed lymphoid panel = 4783 and random seed myeloid panel = 8283. In addition to the unsupervised analytical approach, Manual gating was conducted according to established gating strategies, with slight modifications implemented to improve resolution and allow the identification of additional immune cell populations (17) to validate the findings obtained with unsupervised clustering and to minimise the risk of omitting biologically relevant cell populations (Suppl. Fig. S4) Heatmaps were generated using the figure creation tool within OMIQ. Customised x-axes and coloured continuous features were utilised to assign biological phenotypes to the identified clusters (Suppl. Fig. S6 B,C)

### Statistical analyses

GraphPad Prism version 10.4.1(532) (GraphPad Software, Inc., San Diego, CA, USA) was used for all statistical analyses beyond those described in OMIQ of the FACS data. For non-parametric data, the Kruskal–Wallis test with Dunn’s multiple comparisons post hoc test and a mixed-effects model (REML) were used, followed by Tukey’s multiple comparisons test to assess differences between groups. Longitudinal differences in parametric data sets were analysed with a repeated measures one-way ANOVA followed by Tukey’s post hoc test to correct for multiple comparisons. For analyses involving multiple independent variables, a two-way ANOVA was applied. The results of statistical testing were marked follows: *p < 0.05; **p < 0.01; ***p < 0.001; ****p < 0.0001. A p-value < 0.05 was considered statistically significant. Likelihood ratios (LR) were calculated for each parameter by dividing the percentage of cells of a certain subpopulation found in samples of *de novo* patients by the percentage of cells of that subpopulation found in samples of control negative patients.

## Results

### A representative cohort of AHF patients was recruited to participate in AHF-ImmunoCS

The study cohort had a mean age of 70 years (SD ±12.30 years), reflecting an elderly patient population with typical demographics for heart failure (1): Female patients comprised 36% of the cohort (n = 40). The majority of patients (67%) presented with chronic HF, whereas *de novo* HF was observed in 35 individuals (33%). In terms of aetiology, ischaemic heart disease was the most common underlying cause, identified in 35% of cases, followed by valvular heart disease in 17%. Functional status assessment using the New York Heart Association (NYHA) classification revealed that 58% of patients were in NYHA class III, while 24% were classified as NYHA class IV (Table 1, Suppl. Fig S1, S2).

**Table 1.** Baseline characteristics and laboratory parameters assessed during the index hospitalisation. The study cohort consisted of 110 samples divided into subgroups stratified by the presence of HRA as detected by IFT at baseline (BL) and at the 6-month and 6-week follow-up visits (6W,6M), respectively.

|  | Total<br>N=110 | De novo<br>N=24 | persistently-positive N=52 | persistently-negative N=34 |
| --- | --- | --- | --- | --- |
| Age [years], mean (SD) | 71 (12) | 70 (8) | 70 (14) | 73 (11) |
| Female, n (%) | 40 (36%) | 7(29%) | 18 (34%) | 15 (44%) |
| **HF duration** |  |  |  |  |
| Chronic HF, n (%) | 74 (67%) | 18(75%) | 35 (67%) | 21 (61%) |
| De novo HF, n (%) | 35 (31%) | 6(25%) | 16 (27%) | 13 (38%) |
| **HF type** |  |  |  |  |
| HFrEF, n (%) | 42 (38%) | 10 (41%) | 16 (30%) | 16 (47%) |
| HFmrEF, n (%) | 16 (14%) | 2 (8%) | 12 (23%) | 2 (5%) |
| HFpEF, n (%) | 52 (47%) | 12 (50%) | 24 (46%) | 16 (47%) |
| **Main cause of acute heart failure** |  |  |  |  |
| Valvular heart disease, n (%) | 19 (17%) | 2 (8%) | 9 (17%) | 6 (31%) |
| Arrhythmia, n (%) | 14 (12%) | 0 | 7 (13%) | 4 (21%) |
| Hypertensive heart disease, n (%) | 13 (12%) | 4 (16%) | 3 (5%) | 3 (15%) |
| Ischemic heart disease, n (%) | 35 (32%) | 9 (37%) | 17 (32%) | 4 (21%) |
| Cardiomyopathy, n (%) | 18 (16%) | 7 (29%) | 10 (19%) | 0 |
| Other, n (%) | 11 (10%) | 2 (8%) | 5(10%) | 2 (10%) |
| **NYHA class** |  |  |  |  |
| II, n (%) | 18 (16%) | 4 (22%) | 10 (17%) | 2 (10%) |
| III, n (%) | 64 (58%) | 11 (61%) | 32 (55%) | 10 (52%) |
| IV, n (%) | 27 (24%) | 3 (16%) | 16 (27%) | 7 (36%) |
| **Serum Chemistry** |  |  |  |  |
| Troponin [pg/ml], median (Q1,Q3) | 43.4<br>(24.0/78.1) | 54.4<br>(40.1/68.9) | 32.4<br>(22.4/59.6) | 48.9<br>(26.0,196.4) |
| NT-proBNP [pg/ml], median (Q1,Q3) | 3,791<br>(1,920/11,152) | 3,696<br>(1,982/8,812) | 3,791<br>(1,780/10,163) | 4,085<br>(2,410/20,664) |

### HRA profiling in heart failure patients reveals differences to healthy controls, but no change over time

To characterise HRA expression in HF patients at baseline and 6 weeks as well as 6 months after hospitalisation for AHF, serum antibody levels against the target proteins Troponin I3 (TPI3), Tropomyosin 1 (TPM1), Myosin Heavy Chain 7 (MYH7), Myosin Heavy Chain 6 (MYH6), Myosin Light Chain 7 (MYL7) and the second extracellular loop of the β1-drenergic receptor (β1-EC_II_-AR) we established flow cytometric bead assays (Fig. 1). The antigens were chosen based on the expression patterns reported in the Human Protein Atlas (Suppl. Table 7). Samples of sex- and age-matched donors of the general population without a diagnosis of heart failure (STAAB cohort) (14) were measured in parallel to the samples from HF patients and are referred to as ‘healthy controls’. Compared to healthy controls, AHF patients exhibited elevated antibody levels against TPI3 (Fig. 2A), TPM1 (Fig. 2B) and MYL7 (Fig. 2C) with no significant changes over time from baseline to the six-week and the six-month follow-up. For anti-MYH6, anti-MHY7 and anti-β1-EC_II_-AR antibodies there were no differences in expression levels between AHF patients and healthy controls (Fig. 2D-F). Among the antibody reactivities which were elevated, anti-TPI3 antibody levels were most clearly and consistently increased compared to healthy controls (Fig. 2A) followed by anti-MYL7 (Fig. 2C) and anti-TPM1 antibodies (Fig. 2B). Our data, thus, show raised HRA levels against certain heart-expressed antigens in AHF patients compared to healthy controls, including antibodies against MYL7 found exclusively in the heart (The Human Protein Atlas) (Suppl. Table 7).

**Fig. 2.**
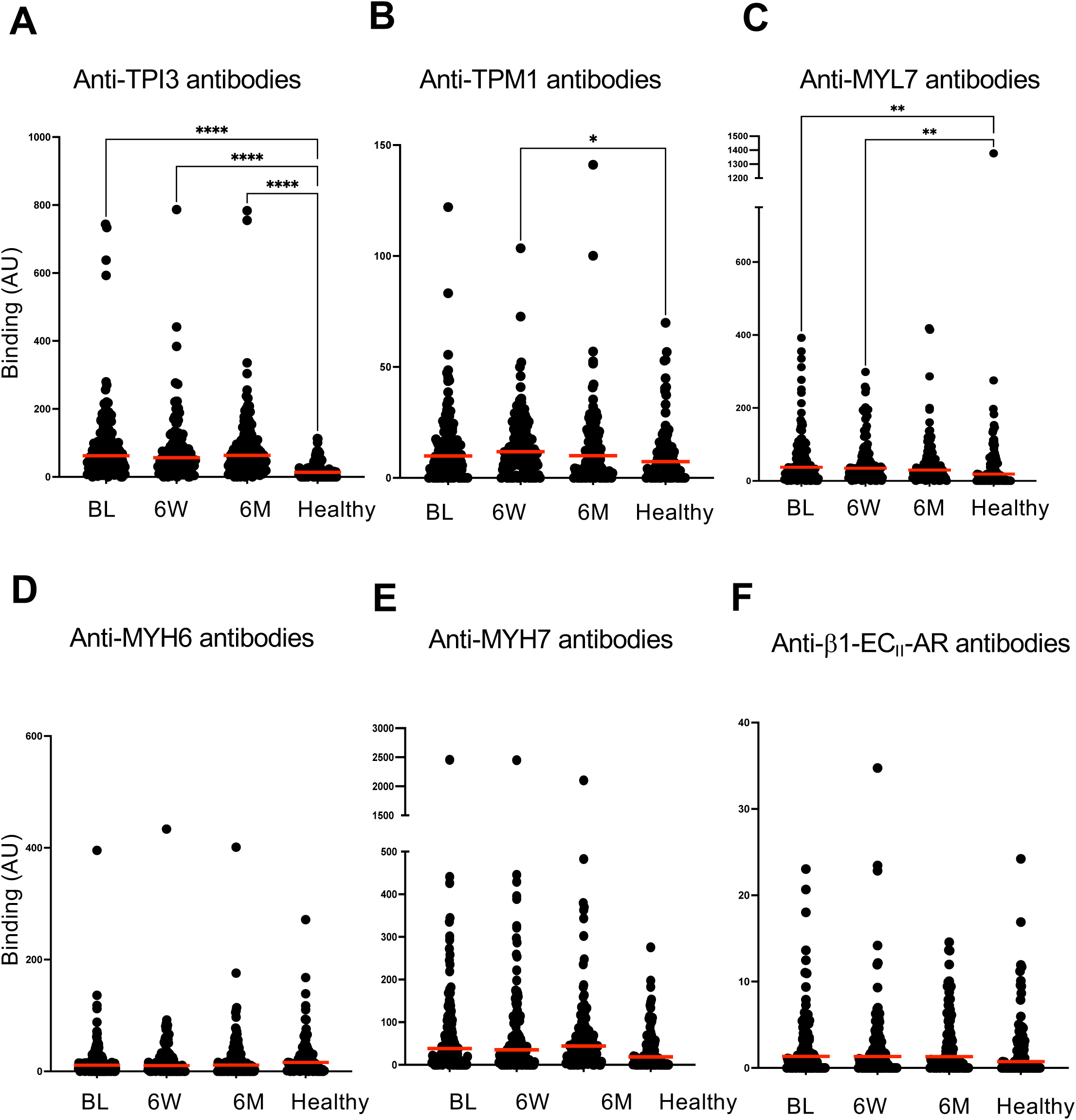
Longitudinal profiling of serum antibody responses in heart failure patients and healthy controls. Serum levels of antibodies targeting (**A**) TPI3, (**B**) TPM1, (**C**) MYL7, (**D**) MYH6, (**E**) MYH7, and (**F**) β_1_-EC_II_-AR were quantified using a bead-based assay in healthy individuals and heart failure patients at baseline, 6 weeks, and 6 months following hospitalization. Results are presented as fluorescence intensity in arbitrary units (AU), which represent the relative level of antibody binding. Statistical analysis was performed using the non-parametric Kruskal–Wallis test with Dunn’s multiple comparisons post hoc test. Statistical significance was defined as follows: *p < 0.05; **p < 0.01; ***p < 0.001; ****p < 0.0001.

### Antibody levels against TPI3, TPM1, MYH7, MYH6, MYL7 and β_1_-EC_II_-AR did not correlate with HRA binding to cardiac tissue as detected by IFT

To further characterise the dynamics of antibody responses against TPI3, TPM1, MYH7, MYH6, MYL7 and β1-EC_II_-AR in relation to IFT status, we first determined binding to the myocardium by IFT and then stratified patients and samples as follows: ‘*De novo’* (patients who were negative in the IFT at baseline and became positive during follow-up), ‘persistently-negative’ (patients with persistently negative samples in the IFT) and ‘persistently-positive’ (patients with persistently positive samples in the IFT, i.e. already at baseline).

Analysis of samples obtained at baseline and 6-month follow-up from the first 110 patients enrolled revealed de novo induction of HRA in 21% of patients, as determined by indirect immunofluorescence testing (IFT), compared with previously published rates of 32% (12).

For the ‘*de novo*’ group we noted that for none of the antibody specificities detected by the bead assays there was a parallel induction of HRA in the IFT (Fig. 3, red circles) meaning that there was no correlation between the antibodies detected by IFT and by bead assay. Similarly, the two other groups, persistently-negative (Fig. 3, grey circles) and persistently-positive (Fig. 3, black circles), also showed no changes over time regarding antibody reactivities measured by bead assay. Between the groups there were also no differences with the exception of antibodies against β_1_-EC_II_-AR which were significantly elevated in the persistently-positive group compared to the *de novo* group (Fig. 3G). Together, we observed that the results of the IFT and the bead assays were independent of each other as there were no correlations between the data obtained with these two test formats.

**Fig. 3.**
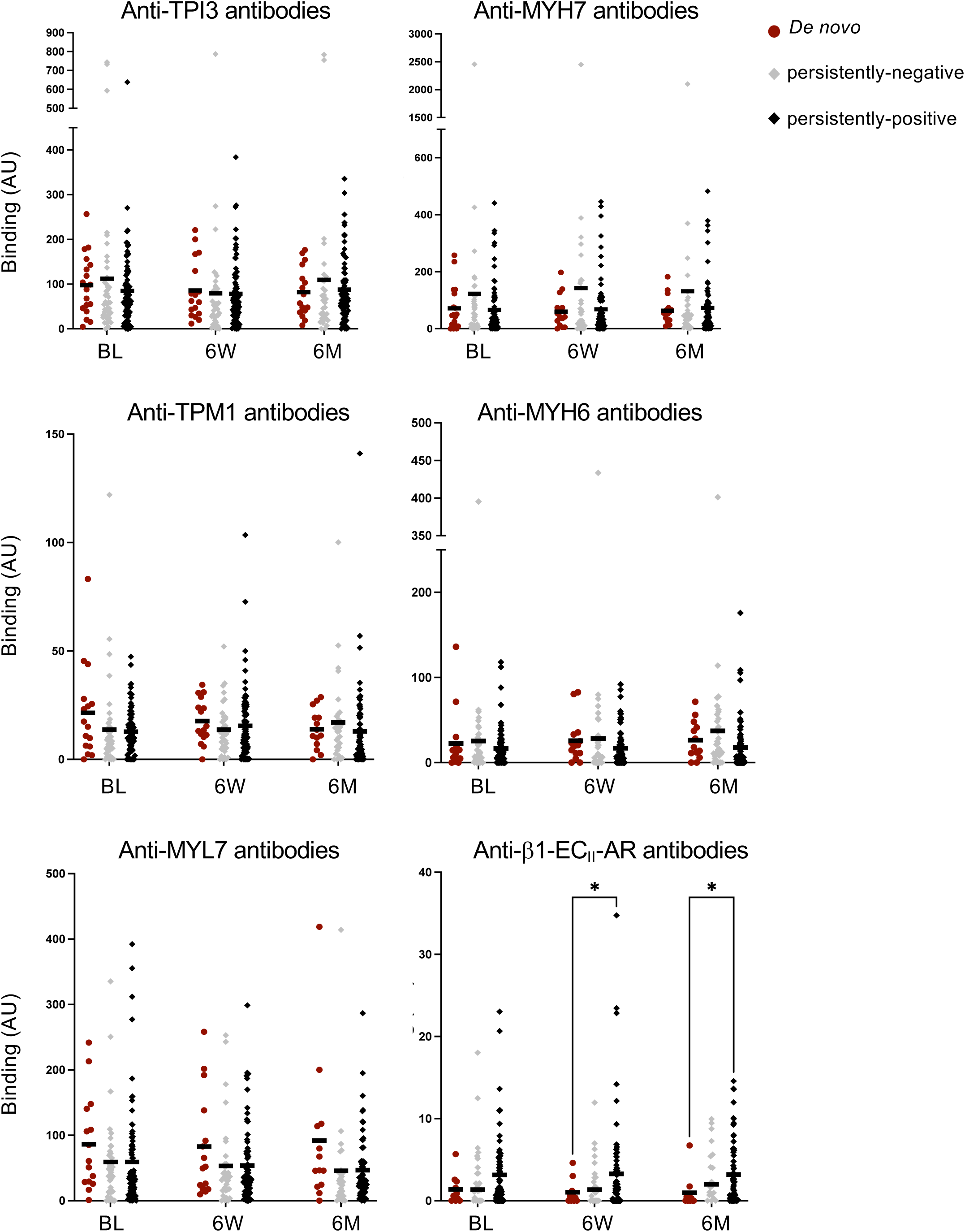
Longitudinal analysis of antibody prevalence against TPI3, TPM1, MYH7, MYH6, MYL7 and β₁_1_-EC_II_-AR stratified by IFT status. Antibody prevalence at baseline, 6 weeks, and 6 months, stratified by IFT classification: *De novo* (patients who became IFT-positive during follow-up), persistently-negative in IFT and persistently-positive in IFT. Statistical analysis was performed using a mixed-effects model (REML), followed by Tukey’s multiple comparisons test to assess differences between groups. Statistical significance was defined as follows: *p < 0.05; **p < 0.01; ***p < 0.001; ****p < 0.0001. For longitudinal analyses only datasets with values for each timepoint were considered and a Friedman test applied rendering no significant differences.

### Differences in the T cell and myeloid cell compartments separated patients stratified according to IFT status

Our autoantibody analyses revealed that AHF did not constitute a trigger for HRA induction across all assays employed by us. Still, in the *de novo* group HRA binding to cardiac tissue is clearly upregulated which might be reflected by changes in the composition of the peripheral immune cell compartment compared to controls. We, thus, analysed thawed whole-blood samples of HF patients that had been obtained at all three time points (baseline, 6 weeks, and 6 months) by high-dimensional flow cytometry (Suppl. Table 1, 2, 3 and Suppl. Fig. S4 - S11). For the identification of leukocyte subpopulations we relied on maker expression due to the well-known changes in the forward/ side scatter profile, particularly of granulocytes, due to freezing and thawing (18, 19). A total of 22 *de novo* patients identified in this study so far were sex- and age-matched to an almost equal number of persistently-negative and persistently-positive patients. We analysed the data by combining unsupervised AI-driven data analysis with manual gating steps to identify well-defined leukocyte subsets (Suppl. Fig. S5). Within the lymphocytic compartment, we observed higher frequencies of CD4⁺ T cells in samples from *de novo* patients compared to control negative patients at baseline and a decrease in *de novo* patients over time (Fig. 4A, Suppl. Fig. S4B). Absolute cell numbers of CD4^+^ T cells per μl of blood did not differ (Fig. 4C) and taking into account the proportion of CD8^+^ T cells among lymphocytes also did not further discriminate the groups (Fig. 4B, D, Suppl. Fig. S4F). Analysis of CD4^+^ T cell subsets revealed higher frequencies of naive conventional, i.e. non-regulatory, CD4^+^ T cells in samples of *de novo* compared to both persistently-negative and persistently-positive patients (Fig. 5A, B, Suppl. Fig. S4B). Subdividing the memory compartment into central and effector memory T cells, again, did not further discriminate the groups (Fig. 5C, D; Suppl. Fig. S4E). In contrast to the reduced proportion of memory cells among CD4^+^ T cells, the proportion of Th1 and a Th17-like cluster (cluster 1008) identified by unsupervised analysis were higher in the *de novo* group than in persistently-positive samples at baseline and partly also at later time points (Fig. 5E, F; Suppl. Fig. S4D, S6D).

**Fig. 4.**
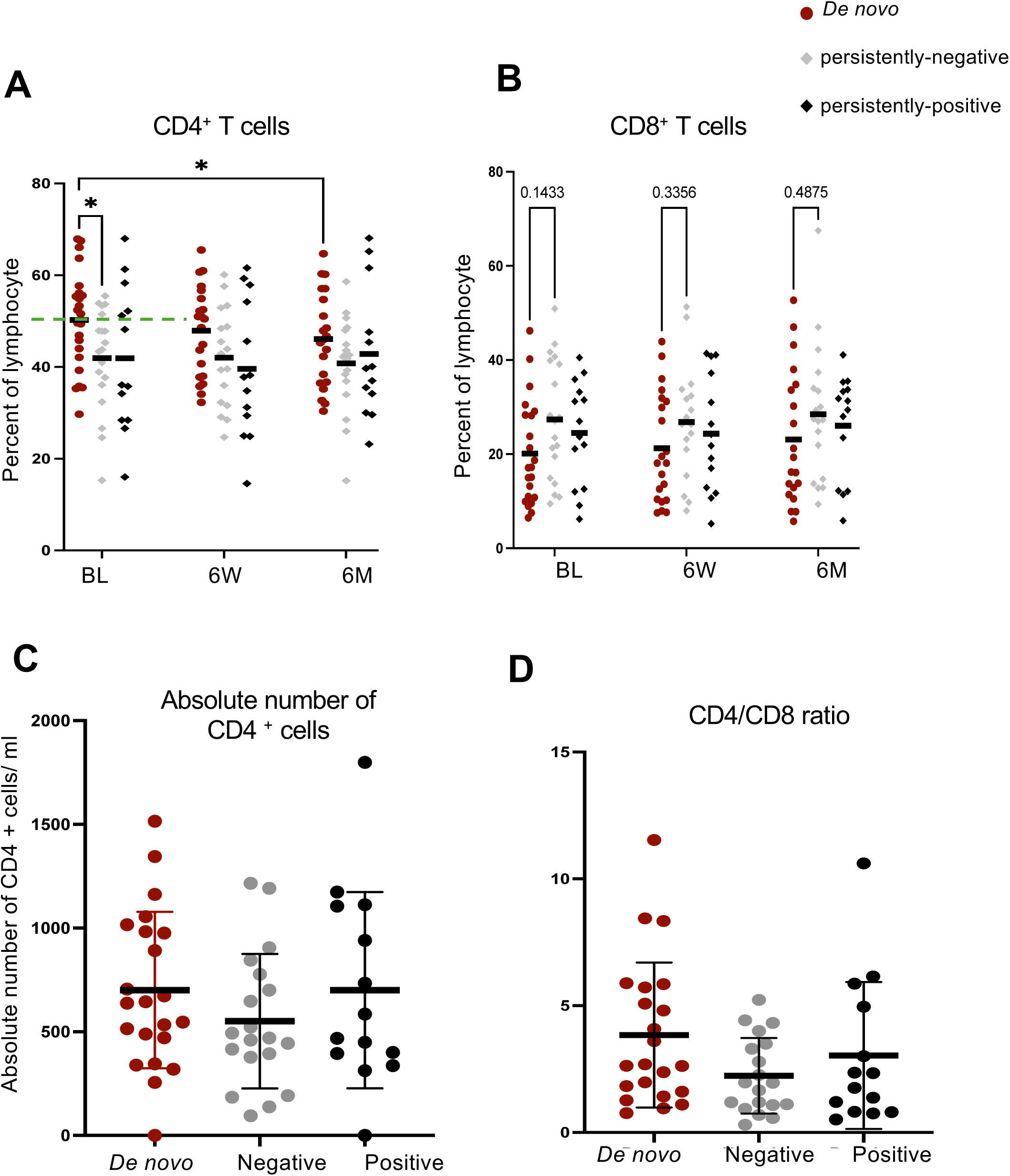
High-dimensional flow cytometry was used to profile peripheral immune cells in heart failure patients across multiple time points (baseline, 6 weeks, and 6 months) in distinct IFT cohorts. (**A**) Frequency of CD4^+^ and (**B**) CD8^+^ T cells among lymphocytes for the three cohorts defined by the IFT results over time. (**C**) Absolute count of CD4^+^ T cells/µ. (**D**) CD4/CD8 ratio. Statistical analysis was performed using the non-parametric Kruskal–Wallis test with Dunn’s multiple comparisons post hoc test. Statistical significance was defined as follows: *p < 0.05; **p < 0.01; ***p < 0.001; ****p < 0.0001.

**Fig. 5.**
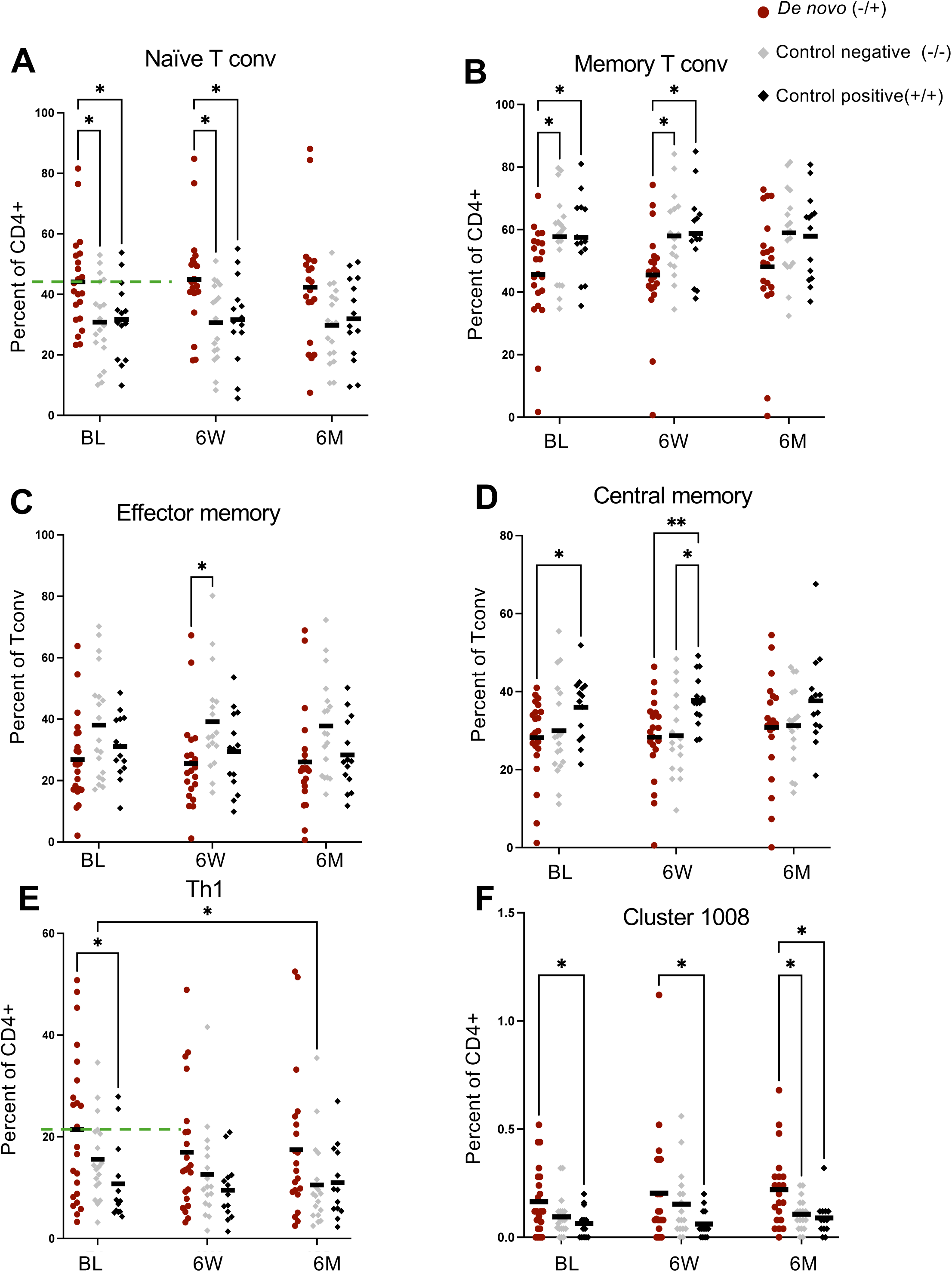
High-dimensional flow cytometry-based profiling of peripheral immune cells in heart failure patients at multiple time points (baseline, 6 weeks, and 6 months) in different IFT cohorts with a focus on CD4^+^ T cell subsampling. Proportion of (**A**) naive Tconv and (**B**) memory Tconv among CD4^+^ T cells. (**C**) Proportion of effector and (**D**) central memory cells among CD4^+^ T conv. (**E**) Percent Th1 and (**F**) Th17-like Cluster 1008 cells among CD4^+^ T cells. Statistical analysis was performed using a mixed-effects model (REML), followed by Tukey’s multiple comparisons test to assess differences between groups. Statistical significance was defined as follows: *p < 0.05; **p < 0.01; ***p < 0.001; ****p < 0.0001.

Among myeloid cells, we also observed differences between the groups in samples taken six weeks and six months after the index hospitalisation. (Fig. 6A-F, Suppl. Fig. S4I-M). At baseline, there were no differences between the groups regarding composition of the myeloid compartment. The overall frequency of total granulocytes did not differ significantly among the study groups (Fig. 6A). At the six-month follow-up, frequencies of eosinophils among myeloid cells were higher for *de novo* versus persistently-positive patients (Fig. 6B). For a subset of mature neutrophils (cluster 28) (Fig. 6C, Suppl. Fig. S6C) and for polymorphnuclear myeloid-derived suppressor cells (PMN-MDSC) (Fig. 6F) *de novo* patients had lower proportions among myeloid cells than persistently-negative and/or persistently-positive patients. Together, our data show that *de novo* samples and the two patient groups with persistent HRA expression could be differentiated based on the composition of the leukocyte compartment with differences regarding CD4^+^ T cells already present at baseline.

**Fig. 6.**
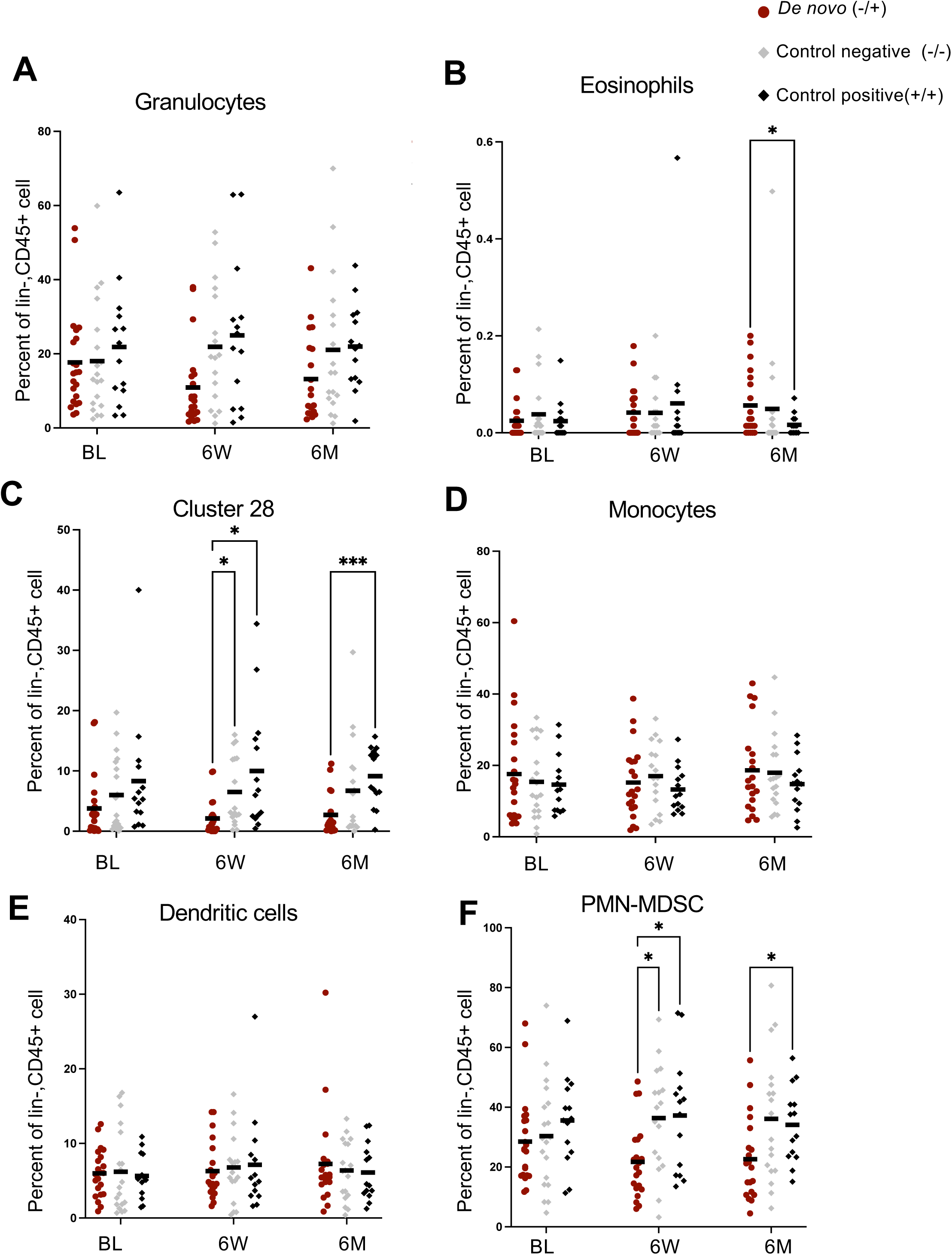
Longitudinal immunophenotyping of the peripheral-blood myeloid compartment in heart failure patients stratified according to IFT categories. Proportion of **(A)** granulocytes, (**B**) eosinophils, (**C**) Cluster 28 mature neutrophils, (**D**) monocytes, (**E**) dendritic cells and (**F**) PMN-MDSC (polymorphonucler myeloid-derived suppressor cells) among myeloid cells. Statistical analysis was performed using a mixed-effects model (REML), followed by Tukey’s multiple comparisons test to assess differences between groups. Statistical significance was defined as follows: *p < 0.05; **p < 0.01; ***p < 0.001; ****p < 0.0001.

### Screening for HRA by IFT plus cellular immunophenotyping allows to prospectively identify patients developing HRA *de novo*

Ideally, it would be possible to define a diagnostic algorithm that allows to identify HRA seroconverters at baseline. According to our pilot data, these patients are at higher risk and might benefit from immune targeting and/or more intensive therapy. For the flow cytometry data, this means optimising cut-offs so that the likelihood ratios for samples from *de novo* over control negative patients are as high as possible - ideally reaching a value around ten (PMID 38300842). Therefore, we calculated the likelihood ratios for the proportion of CD4^+^ T cells among lymphocytes, the percentage of naive Tconv and Th1 cells among CD4^+^ T cells comparing *de novo* and control negative samples (Suppl. Fig. S12). The cut-offs yielding the highest likelihood ratios, while still providing for statistical robustness (at least 10% per group above the threshold), are shown as green dashed lines in Fig. 4A, 5A, E). The resulting LR values for the three T cell subsets analysed were 2.1, 3.0 and 3.3, respectively. Moreover, for each T cell subset, values above a certain threshold were either exclusive or almost exclusive to HRA seroconverters (Suppl. Fig. S12, pink dashed line). The data, thus, imply that it might be possible to identify HRA seroconverters already at baseline by combining autoantibody testing by IFT with cellular immunophenotyping analysing the proportion of CD4^+^ T cells among lymphocytes as well as the frequencies of naive Tconv and Th1 cells among CD4^+^ T cells.

## Discussion

In this study we followed up on our previous observation that induction of HRA in the wake of acute decompensation of HF was associated with a poor clinical prognosis (12). Using a panel of assays detecting HRA against defined antigens expressed in the heart, we, however, found in this study that AHF did not broadly induce the generation of autoantibodies against defined cardiac antigens. But we noted that HRA levels e.g. against TPI3 were already higher in HF patients at baseline compared to healthy controls, potentially indicating that cardiac damage originating from the underlying disease or due to chronic HF induced these higher HRA levels. Cellular immunophenotyping further revealed that changes in the composition of peripheral blood leukocytes, particularly T cells, preceded induction of HRA, potentially positioning T cells high up in the pathophysiological cascade leading to HRA induction.

As B cells are *per se* highly tolerant towards intracellular antigens (20), it is mainly down to CD4^+^ T cells to control the induction of autoantibodies. ‘Holes’ in central T cell tolerance as a result of thymic selection, thus, directly impact autoantibody induction, as is the case for MYH6 (α-MyHC), which is not expressed in the thymus (21).

Generally, autoantibodies against cardiac antigens are believed to either be disease-inducing, as is the case for anti-β1-EC_II_-AR antibodies (10, 11, 22), or to exacerbate myocardial injury, as is the case e.g. for troponin I-specific antibodies (reviewed in (23)). However, our observation that HRA against cardiac antigens can also be found in healthy individuals (Fig. 2), where they are associated with a favourable cardiovascular risk profile and better cardiac function (unpublished data), implies that HRA might also exert cardioprotective effects.

We used an immunofluorescence test with heart muscle tissue as the substrate to identify the relationship of HRA induction and clinical prognosis (12). To obtain a broader picture of the autoantibody response induced against cardiac antigens, we set up a number of bead- and flow cytometry-based assays to measure these antibodies. However, none of the antibodies detected by one of the bead-based assays correlated with the results of the IFT (Fig. 1, 3) and there was also no increase in the reactivity in the bead-based assays over time (Fig. 2). This also holds true for antibody affinity, for which we also saw no increase from baseline to the follow-up time points (unpublished data). Currently, it is unclear whether any of the autoantibodies detected by bead assay are associated with clinical prognosis to a similar or even stronger degree than the autoantibodies generating a striated pattern in the immunofluorescence test. However, analysis of the data generated in AHF-ImmunoCS upon completion of the study would ideally confirm our initial observation using the immunofluorescence test (12) and reveal further associations of autoantibody expression with severity and prognosis of AHF.

Clinically and with respect to the concentration of clinical chemistry parameters such as NT-proBNP, the data of this cohort are in line with our previous study (12), showing that, at baseline, HF in patients developing HRA is not more severe than in the other HF patients (Table 1). This corroborates that HRA constitute an independent bio- and prognostic marker, i.e. HRA levels do not simply mirror the stage of the disease and the level of standard clinical chemistry parameters at baseline.

To understand whether changes in the composition of peripheral blood leukocytes are associated with the induction of HRA and to gain insights into the underlying pathomechanism of HRA induction, we immunophenotyped both the lymphoid and the myeloid compartments in peripheral blood using spectral flow cytometry. The combination of manual gating and unsupervised clustering allowed us to identify phenotypes that are associated with the induction of HRA, with some phenotypes already detected at baseline (Fig. 4 - 6). Regarding the latter, the CD4^+^ T cell compartment stuck out, suggesting that the higher proportion of CD4^+^ T cells among lymphocytes and the higher percentage of Th1 cells among CD4^+^ T cells might indicate that in these patients, B cells receive more ‘help’ from CD4^+^ T cells with Th1-derived IFNγ driving antibody class-switching to IgG (24). For CD4^+^ T cells we did not observe an increase in the activation status (markers: CD38, HLA-DR) or proliferation of the cells (marker: Ki-67) (unpublished data). This means that activation of the cells might primarily happen in lymph nodes or the spleen. The higher percentage of naive cells among conventional CD4^+^ T cells in patients who go on to develop HRA *de novo* might reflect preferential retention of memory CD4^+^ T cells in the tissue. Particularly, as the total number of CD4^+^ T cells was not increased in peripheral blood (Fig. 4). A similar consideration might also apply to B cells, for which we could not detect any differences in the peripheral blood between samples from the HRH *de novo* group and the other two patient cohort defined by HRA status in the IFT (Suppl. Fig. S9).

In our analyses, we did not characterise T and/ or B cells with specificity for cardiac antigens. It may, thus, be that analysing the prevalence and activation status of cardiac antigen-specific B and T cells in peripheral blood might show differences between the groups. The sheer presence of cardiac antigen-specific B cells in peripheral blood would indicate activation of these cells as has been shown for B cells reacting to a booster vaccination (25). However, for vaccine antigen-specific B cells the time window during which the cells could be detected in peripheral blood was roughly only about a day, complicating their detection (25).

While CD4^+^ T helper cells are perfectly suited to enhance HRA induction, PMN-MDSC are well known for their immunosuppressive properties from studies into cancer and autoimmunity (reviewed in (26–29). A lower proportion of PMN-MDSC among myeloid cells in the HRA *de novo* group versus the two control groups became apparent at the six-week follow-up time point (Fig. 6). Lower frequencies of PMN-MDSC among myeloid cells in peripheral blood mean that there was a reduced capacity of these cells to mediate immunosuppression, which might have contributed to HRA induction. Apart from their numbers, the per-cell activity is, of course, also important for the overall suppressive capacity of PMN-MDSC. Here, catecholamines play an important role as they enhance MDSC function by signalling through the β2-adrenergic receptor expressed by these cells (30). Due to the detrimental effects of excessive catecholamine levels on cardiac function, β-blockers are recommended for many HF patients (1). As all of the drugs available not only block the β1, but also the β2 adrenergic receptor to varying degrees (31), HF patients with a high risk of developing HRA *de novo* might profit from receiving the most β1-selective β-blockers and/or from a transiently lower dose of β-blocker to allow for better MDSC activation after the AHF episode.

Regarding the pharmacotherapy of HF, recent years have brought about a significant change with the advent of sodium-glucose cotransporter-2 (SGLT2) inhibitors (32–36). We, thus, hypothesised that the slightly lower percentage of patients developing HRA *de novo* in this study compared to our previous study might be related to SGLT2 inhibitor treatment, given their immunomodulatory properties (reviewed in (37)). However, in our cohort there were no differences in SGLT2 inhibitor use between patients developing HRA *de novo* and either of the two other patient groups (Suppl. Fig. S2B).

Taken together, our study highlights that *de novo* induction of HRA - as detected by immunofluorescence on heart tissue slides - is preceded by changes in the CD4^+^ T cell compartment, which in the future might help to identify these patients before they progress to developing HRA. Moreover, a putative lack of PMN-MDSC might, on the one hand, contribute to HRA induction and, on the other hand, hold the key to novel tailored therapeutic regimens aiming at preventing HRA induction. Upon completion of AHF-ImmunoCS, we will be able to verify the findings put forward here and correlate our biomarker data with clinical outcome.

## List of abbreviations

β1-EC_II_-AR: second extracellular loop of the β1 adrenergic receptor
HRA: heart-reactive antibodies
IFT: immunofluorescence test on cardiac tissue
MYH6: Myosin Heavy Chain 6
MYH7: Myosin Heavy Chain 7
MYL7: Myosin Light Chain 7
PBS: phosphate-buffered saline
PMN-MDSC: polymorphnuclear myeloid-derived suppressor cells
TPM1: Tropomyosin 1
TPI3: Troponin I3

## Acknowledgements

The authors are very grateful to Daria Grosser for her work as a lab assistant measuring and analysing sera of healthy controls. Moreover, the authors like to thank Monika Hanke for expert data management, Daniela Vilsmaier for managing clinical visits and sample collection as well as the Interdisciplinary Bank of Data and Biomaterials Würzburg (ibdw) for storage of whole-blood and serum samples.

