## Supplemental Material for "Changes in peripheral blood leukocyte composition precede development of heart-reactive autoantibodies in patients hospitalised for acute heart failure"

* equal contribution

#### Supplemental Material

**Suppl. Fig. S1. Cardiac biomarker levels and clinical classification of heart failure across IFT cohorts.** (**A**) Troponin levels (ng/l), (**B**) NT-proBNP, N-terminal prohormone of brain natriuretic peptide, levels (pg/ml) and (**C**) distribution of heart failure severity based on New York Heart Association (NYHA) classification are shown for the different IFT cohorts.

**Suppl. Fig. S2. Clinical characteristics and treatment distribution across IFT cohorts.** (**A**) Distribution of heart failure phenotypes across the IFT cohorts. (**B**) Proportion of patients in each IFT cohort receiving SGLT2 inhibitors; this analysis was performed to assess potential associations with the frequency of the *De novo* group. (**C**) Pie chart illustrating the gender distribution within the study cohort.

**Suppl. Fig. S3. Schematic overview of the bead-based antibody detection assay.** Beads were coated with recombinant proteins of interest. Upon incubation with patient samples, antibodies specific for the coated antigens — if present — bind to the corresponding protein-coated beads. Detection was performed using a fluorophore-conjugated secondary antibody targeting human IgG. To validate assay performance, a monoclonal antibody was used as a positive control and AB serum served as a negative control. Samples were acquired on a FACS Calibur flow cytometer.

**Suppl. Fig. S4. Gating strategy for flow cytometric immune phenotyping.** This figure illustrates the gating strategy applied to flow cytometry data from both antibody panels. Panel 1 was used for lymphoid cell identification and Panel 2 for myeloid cell characterisation. Gating was performed in parallel with unsupervised clustering analyses to ensure robust identification of immune cell subsets. Please note that due to freezing and thawing most of the granulocytes ($CD45^{low}$) were $SSC^{low}$.

**Suppl. Fig. S5. Comparison of manual gating and unsupervised clustering across immune cell subsets.** This figure shows side-by-side comparisons of manual gating and unsupervised clustering results for different immune cell populations. (**A**) All lymphocytes, (**B**) all myeloid, (**C**) $CD4^+$ T cell subsampling, (**D**) $CD8^+$ T cell subsampling, (**E**) B cell subsampling and (**F**) NK cell subsampling are depicted. Both approaches were applied to flow cytometry data from Panel 1and 2 to evaluate consistency in immune cell identification.

**Suppl. Fig. S6. Different visualisation strategies in the OMIQ platform used to annotate cell populations.** (**A**) Scatter plots of various marker expressions illustrate the levels and distribution of marker expression across the dataset, aiding in the interpretation of population-specific profiles. (**B**) A heatmap provides an overview of marker expression across clusters, further supporting the identification and annotation of distinct immune cell populations. (**C**) As an example, a specialised X-axis plot highlights that Cluster 28 within the myeloid compartment can be identified as mature neutrophils, while Cluster 8 from a subsampled $CD4^+$ T cell population shows characteristics of Th17-like cells, based on their marker expression patterns.

**Suppl. Fig. S7. Comparative abundance of immune cell types by IFT group at baseline (BL).** The Heatmap highlights differences in immune cell population frequencies across groups, with particular attention to cell types that enable discrimination of the *De novo* group from other control groups at baseline. These distinct immune signatures may serve as potential early indicators of disease onset or group-specific immune states.

**Suppl. Fig. S8. Immune cell signatures analysed over time across IFT groups.** This figure shows the temporal dynamics of immune cell populations, highlighting changes in cell type abundance and phenotype across different IFT groups. Longitudinal analysis allows for the identification of group-specific immune trajectories and potential biomarkers associated with disease progression or resolution.

**Suppl. Fig. S9. Temporal dynamics of B cell populations across IFT groups.** The figure illustrates changes in B cell subset frequencies and phenotypes over time within different IFT-defined groups.

**Suppl. Fig. S10. Frequency of $CD8^+$ and γδ T cell subtypes in IFT-defined groups.** The figure shows the distribution of various $CD8^+$ T cell subsets and natural killer–like γδ T cell populations among participants categorised according to IFT results

**Suppl. Fig. S11. Temporal dynamics of myeloid cell populations across IFT groups.** The figure depicts changes in the frequency and phenotype of neutrophils over time within IFT-defined groups.

**Suppl. Fig. S12. The table presents the likelihood ratios for $CD4^+$ T cells, Th1 cells and naïve $CD4^+$ conventional T (Tconv) cells based on the depicted cut-off lines.**

**Suppl. Table 1. Baseline characteristics and laboratory parameters assessed during the index hospitalisation.** The subcohort used for immunphenotyping consisted of a total of 55 divided into three subgroups based on the presence of HRA at baseline (BL) and at the 6-month and 6-week follow-up visit (6W,6M), respectively.

**Suppl. Table 2. Antibody panel used for immunophenotyping the lymphoid compartment.**

**Suppl. Table 3. Antibody panel used for immunophenotyping the myeloid compartment.**

**Suppl. Table 4. Overview of the proteins and peptides used for the detection of HRA by bead assay.**

**Suppl. Table 5. Summary of the primary antibodies used in every experimental run to verify sufficient coating of beads.**

**Suppl. Table 6. List of secondary antibodies applied for detection in the bead-based assays.**

**Suppl. Table 7. Expression of autoantigens in heart muscle and other tissues according to the Human Protein Atlas.**

### Suppl. Fig. S1

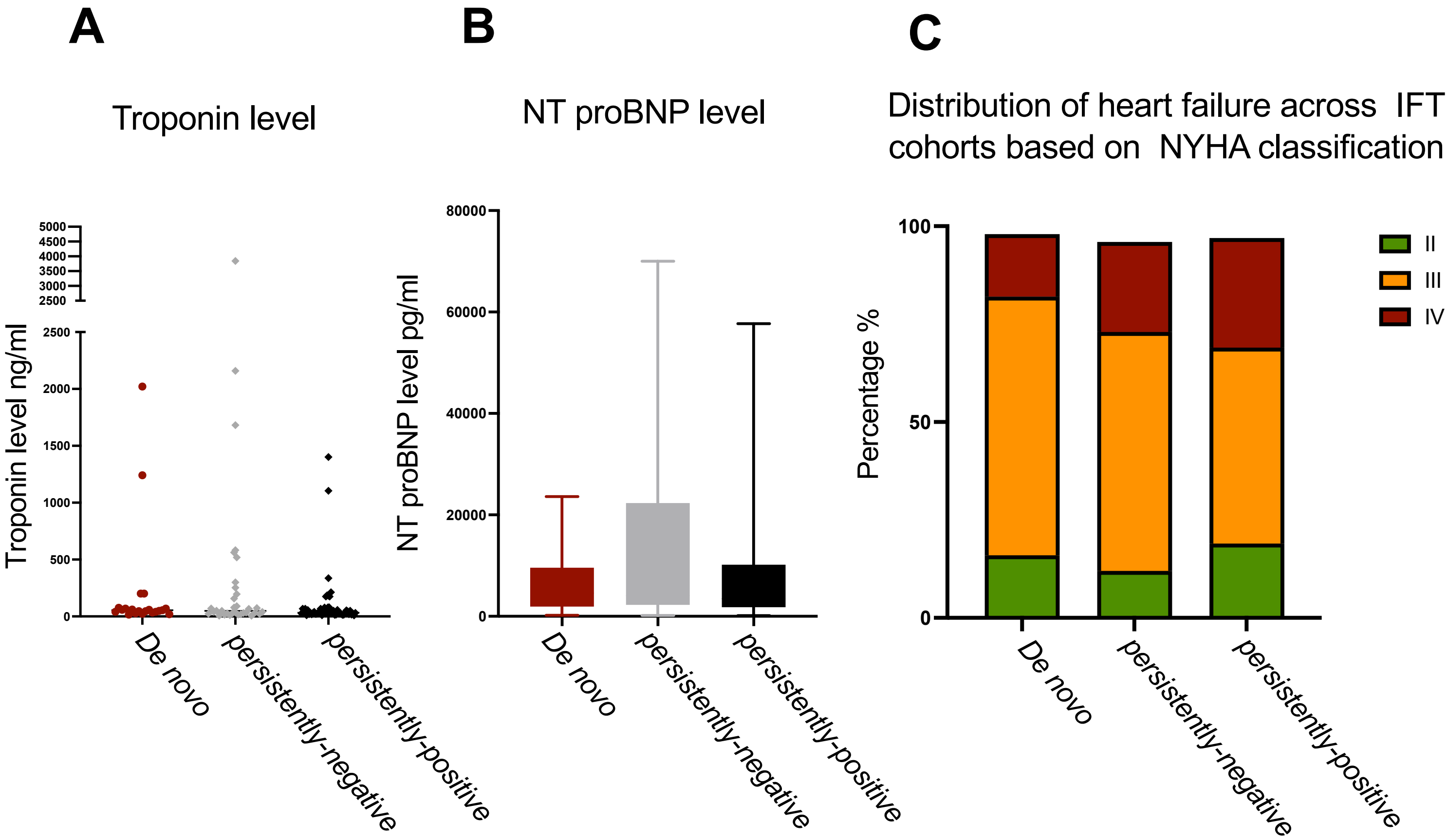


### Suppl. Fig. S2

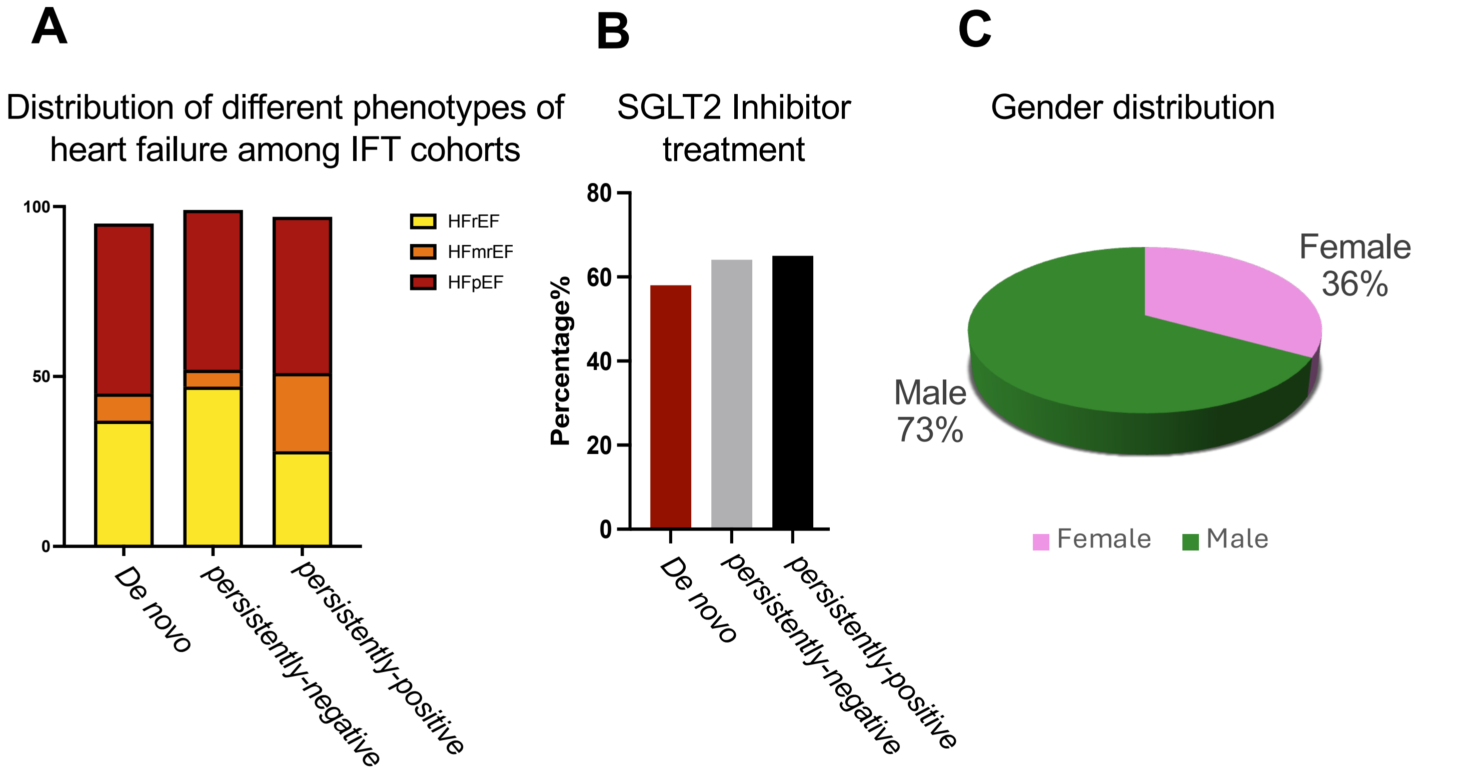

#### Suppl. Fig. S3

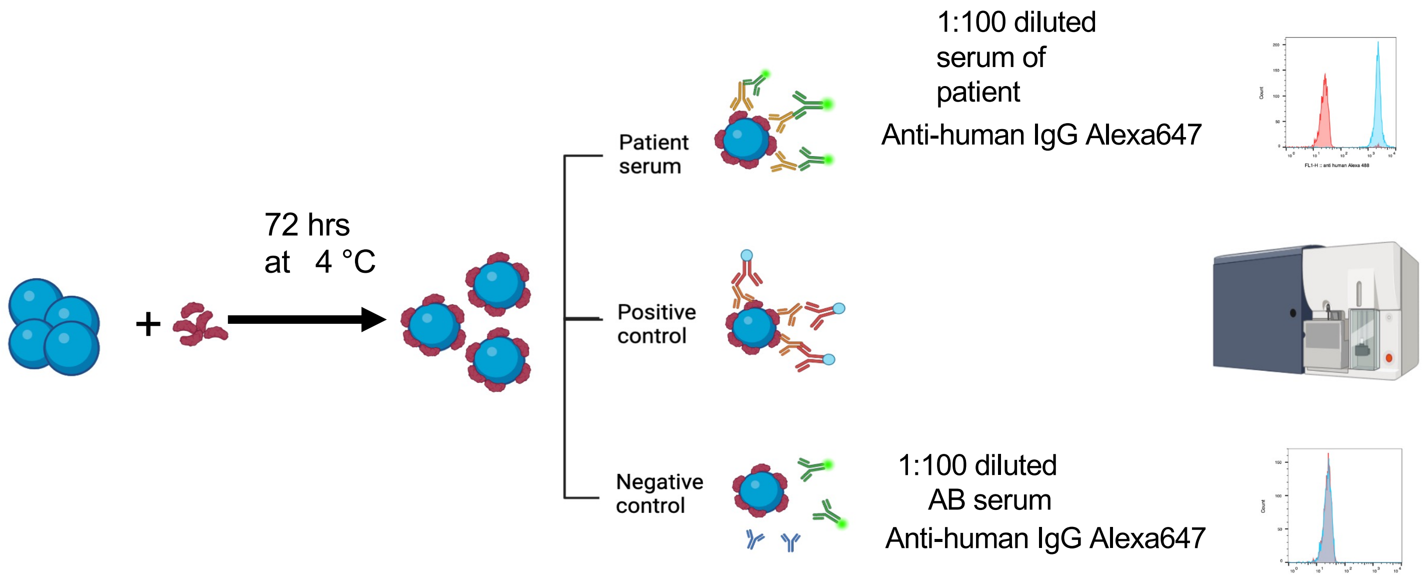

72 hrs
at 4 °C
Patient serum
Positive control
Negative control
1:100 diluted serum of patient
Anti-human IgG Alexa647
1:100 diluted AB serum
Anti-human IgG Alexa647

### Suppl. Table 1

| Baseline characteristics | Total N=55 | Denovo N=22 | persistently-positive N=14 | persistently-negative N=19 |
|---|---|---|---|---|
| **Age** | 74 (37;91) | 72 (55;88) | 71 (37;88) | 80.5 (52;91) |
| **Female** | 15(27%) | 6(33%) | 3(21%) | 6(31%) |
| **Male** | 40(72%) | 16(72%) | 11(78%) | 13(68%) |
| **Chronic** | 37(67%) | 16(72%) | 9(64%) | 12(63%) |
| **Denovo (new)** | 18(32%) | 6(27%) | 5(35%) | 7(36%) |
| **HFrEF n, (%)** | 21(38%) | 10 (45%) | 4(28%) | 8 (42%) |
| **HFmrEF n, (%)** | 6 (10%) | 2 (9%) | 3 (21%) | 1 (5%) |
| **HFpEF n, (%)** | 27 (49%) | 10 (45%) | 7 (50%) | 10 (52%) |
| **Main cause of acute heart failure Heart valve disease n, (%)** | 8 (14%) | 2 (9%) | 0 | 6 (31%) |
| **Heart arrhytmia n, (%)** | 7 (13%) | 0 | 3 (21%) | 4 (21%) |
| **Hypertensive Heart disease n, (%)** | 7 (13%) | 4 (18%) | 1 (5.6%) | 3 (15%) |
| **Ischmisc causes n, (%)** | 18 (32%) | 8 (36%) | 6 (42%) | 4 (21%) |
| **Cardiomyopathy n, (%)** | 9 (16%) | 7 (31%) | 2 (14%) | 0 |
| **Others n, (%)** | 6 (10%) | 1 (5%) | 2 (14%) | 2 (10%) |
| **NYHA II n, (%)** | 12 (21%) | 4 (18%) | 6 (42%) | 2 (10%) |
| **NYHA III n, (%)** | 28 (50%) | 14 (63%) | 4 (28%) | 10 (52%) |
| **NYHA IV n, (%)** | 15 (27%) | 4 (18%) | 4 (28%) | 7 (36%) |
| **NT-proBNP (Median) n, (%)** | 3447 | 4551 | 2846.5 | 4081 |

### Suppl. Table 2

| | Marker | Fluorophore | Clone | Catalog number | Supplier |
|---|---|---|---|---|---|
| 2 | CD45 RA | BUV395 | 5H9 | 740315 | BD |
| 3 | CD4 | BUV496 | SK3 | 612936 | BD |
| 4 | CD24 | BUV563 | ML5 | 741364 | BD |
| 5 | CD19 | BUV661 | HIB19 | 741604 | BD |
| 6 | Tim-3 | BUV737 | 7D3 | 748820 | BD |
| 7 | CD8 | BUV805 | SK1 | 612889 | BD |
| 8 | CCR7 | BV421 | G043H7 | 353208 | Biolegend |
| 9 | CD16 | SB436 | 3G8 | 62-0166-42 | Thermofisher/eBio science |
| 10 | NKG2a | Pac Blue | S19004C | 375110 | Biolegend |
| 11 | TIGIT | BV480 | 741182 | 747843 | BD |
| 12 | CD3 | BV510 | SK7 | 344828 | biolegend |
| 13 | HLA-DR | BV570 | L243 | 307638 | Biolegend |
| 14 | CD28 | BV605 | CD28.2 | 302968 | Biolegend |
| 15 | CCR4 | BV650 | 1G1 | 744140 | BD |
| 16 | yd TCR | BV711 | 11F2 | 745505 | BD |
| 17 | PD-1 | BV750 | EH12.2H7 | 329966 | biolegend |
| 18 | CTLA-4* | BV785 | BNI3 | 369624 | Biolegend |
| 19 | CCR6 | BB515 | 11A9 | 564479 | BD |
| 20 | CD56 | Alexa488 | HCD56 | 318312 | Biolegend |
| 21 | CD20 | Alexa 532 | 2H7 | 58-0209-42 | Thermofisher/eBio science |
| 22 | CD45 | PerCP | 2D1 | 368506 | Biolegend |
| 23 | Ki67* | PerCP-eFluor710 | SolA15 | 46-5698-82 | Thermofisher/eBio science |
| 24 | KLRG1 | PE | 2F1/KLRG1 | 138408 | Biolegend |
| 25 | CXCR5 | PE-Dazzle | J252D4 | 356928 | Biolegend |
| 26 | CD25 | Spark NIR685 | M-A251 | 356152 | Biolegend |
| 27 | CXCR3 | PE-Cy7 | G025H7 | 353720 | Biolegend |
| 28 | IgD | APC | IA6-2 | 348222 | Biolegend |
| 29 | CD15s | A647 | FH6 | 368103 | Biolegend |
| 30 | Foxp3* | PE-Cy5.5 | | | Thermofisher |
| 31 | CD27 | APC-H7 | M-T271 | 560222 | BD |
| 32 | CD38 | APC-Fire 810 | HIT2 | 356643 | BioLegend |
| 33 | CX3CR1 | BUV615 | 2A9-1 | 751513 | BD |
| 34 | CD62L | R718 | DREG-56 | 567988 | BD |

### Suppl. Table 3

| | Marker | Fluorophore | Clone | Catalog number | Supplier |
|---|---|---|---|---|---|
| **1** | Siglec8 | BUV 395 | 837535 | 747876 | BD |
| 2 | CD33 | BUV496 | WM53 | 741148 | BD |
| 3 | CD226 | BUV563 | DX11 | 748429 | BD |
| 4 | CD69 | BUV661 | FN50 | 750213 | BD |
| 5 | CD155 | BUV737 | SKII.4 | 748586 | BD |
| 6 | CD64 | BUV805 | 10-Jan | 742023 | BD |
| 7 | Tim-3 (CD366) | BV421 | 7D3 | 565562 | BD |
| 8 | CD123 | SB436 | 6H6 | 62-1239-42 | Thermofisher |
| 9 | CD15 | Pacific Blue | W6D3 | 323022 | Biolegend |
| 10 | CD86 | BV480 | 2331 | 566131 | BD |
| 11 | CD16 | BV570 | 3G8 | 302036 | Biolegend |
| 12 | CD83 | BV650 | HB15e | 740602 | BD |
| 13 | CD34 | BV711 | 8G12 | 745543 | BD |
| 14 | CD14 | BV750 | 63D3 | 367136 | Biolegend |
| 15 | CD303 (BDCA2) | BV785 | 201A | 354222 | Biolegend |
| 16 | CD141 (BDCA3) | BB15 | 1A4 | 565084 | BD |
| 17 | CD56 | A488 | HCD56 | 318312 | Biolegend |
| 18 | CD3 | 488 | SK7 | 344810 | biolegend |
| 19 | CD19 | 488 | HIB19 | 302219 | Biolegend |
| 20 | CD11c | A532 | Klon 3.9 | 58-0116-41 | Thermofisher |
| 21 | CD45 | PerCp | 2D1 | 368506 | Biolegend |
| 22 | RANK (CD265) | PE | R12-31 | 119806 | Biolegend |
| 23 | PDL-1 | PE-Dazzle 594 | 29E.2A3 | 329732 | Biolegend |
| 24 | CD11b | PE-AlexaFluor610 | M1/70 | 61-0112-82 | Thermofisher |
| 25 | CXCR3 | PE-Cy7 | G025H7 | 353720 | Biolegend |
| 26 | CD1c (BDCA1) | A647 | L161 | 331510 | Biolegend |
| 27 | CD62L | A700 | DREG-56 | 304820 | Biolegend |
| 28 | CD66b | APC-Cy7 | G10F5 | 305126 | Biolegend |
| 29 | HLA-DR | APC-Rire 810 | L243 | 307674 | Biolegend |
| 30 | | | | | |
| 31 | Ki67 | BV510 | Ki67 | 350518 | Biolegend |
| 32 | Phospho-S6 | PerCp-eFluor710 | cupk43k | 46-9007-42 | Thermofisher |
| 33 | Arg1 | APC | 14D2C43 | 369706 | Biolegend |

#### Suppl. Table 4

| **Protein/ peptide** | **Molecular weight** (kDa) | **Sequence** | **Catalog number** | **Manufacturer** |
|---|---|---|---|---|
| Troponin I3 | 24 | Full length | T9924 | SIGMA-ALDRICH |
| Tropomyosin | 35 | Full length | Pro-469 | PROSPEC |
| MYH7 | 62 | Full length | RPP164Hu01 | Cloud-Clone Crop |
| MYH6 | 223.6 | Full length | TP313673 | ORIGENE |
| MYL7 | 22 | Full length | ab126669 | Abcam |
| Beta1-EC$_{II}$-AR Linear peptide | 2.94 | H-ARAESDEARR CYN DPKCCDFVTN RQ-OH | - | Peptide Specialty Laboratories GmbH |
| Beta1-EC$_{II}$-AR Scrambled peptide | 2.94 | H-SFAVRDERA CRKY QT ACDDCNRNE P-OH | - | Peptide Specialty Laboratories GmbH |

#### Suppl.Table 5

| Antigen | Clone | isotype | species | Catalog number | Manufacturer | Dilution |
|---|---|---|---|---|---|---|
| Troponin I3 | Polyclonal | IgG | Rabbit | 21652-1-AP | Proteintech | 1:1000 |
| Tropomyosin | Polyclonal | IgG | Rabbit | MBS824770 | MyBioSource | 1:1000 |
| 6x His Tag (C-Term)-FITC | Monoclonal 3D5 | IgG2b | Mouse | R933-25 | Invitrogen | 1:500 |
| C-Myc Tag | Monoclonal 4A6 | IgG1 | Mouse | 05-724-25UG | Sigma-Aldrich | 1:1000 |
| MYL7 | Polyclonal | IgG | Rabbit | ab205374 | Abcam | 1:1000 |
| Beta1-$EC_{II}$-AR Linear peptide | Monoclonal 13F6 | | Rat | | In house production | 1:115 |

#### Suppl.Table 6

| Antigen | Fluorophore | clone | Catalog number | manufacturer |
|---|---|---|---|---|
| Human IgG (H+L) Affini pure TM | Alexa Flour 647 | Polyclonal | 109-605-088 | Jackson ImmunoResearch Laboratories |
| Mouse IgG (H+L) Affini pure TM | FITC | Polyclonal | 115-095-166 | Jackson ImmunoResearch Laboratories |
| Rabbit IgG (H+L) Affini pure TM | Alexa Flour 488 | Polyclonal | 111-545-045 | Jackson ImmunoResearch Laboratories |
| Rat IgG (H+L) | PE | Polyclonal | PN IM1622 | Beckman Coulter |

#### Suppl. Table 7

| Protein | Skeletal muscle | Smooth muscle | Heart muscle | Other tissues |
|---|---|---|---|---|
| MYH 7 | ✓ | ✗ | ✓ | ✗ |
| MYH 6 | ✓ | ✗ | ✓ | ✗ |
| MYL 7 | ✗ | ✗ | ✓ | ✗ |
| Tropomyosin 1 | ✓ | ✓ | ✓ | ✓ Appendix, Testis,Intestine, Oral Mucosa |
| Troponin I3 | ✗ | ✗ | ✓ | ✓ Testis |
| β1 adrenergic receptor | ✗ | ✗ | ✓ | ✓ |

THE HUMAN PROTEIN ATLAS

### Suppl. Fig. S4

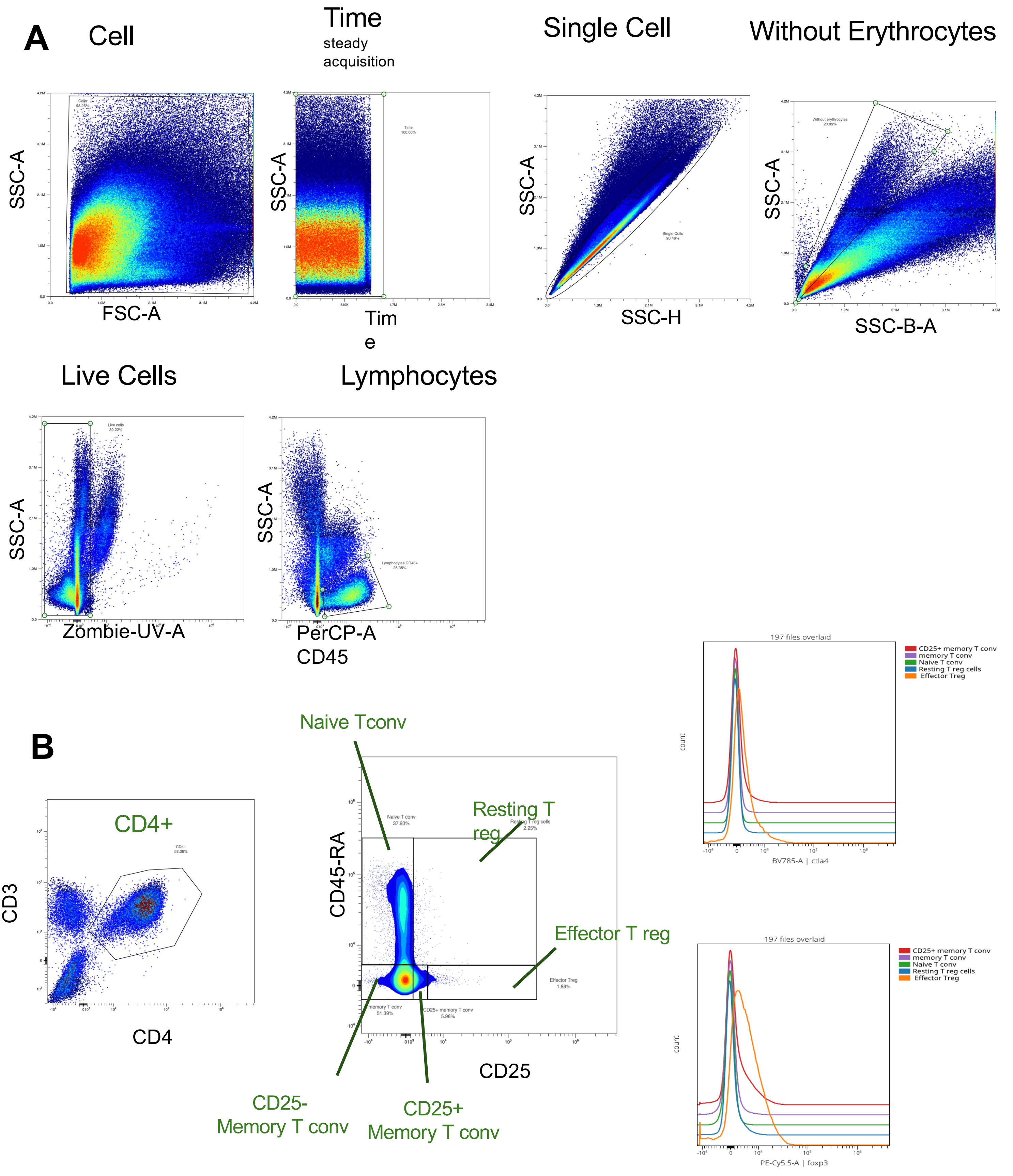
A
Cell
Time
steady acquisition
Single Cell
Without Erythrocytes
SSC-A
FSC-A
Time
SSC-H
SSC-B-A
Live Cells
Lymphocytes
Zombie-UV-A
PerCP-A CD45
B
CD4+
CD3
CD4
Naive Tconv
Resting T reg
Effector T reg
CD45-RA
CD25
CD25- Memory T conv
CD25+ Memory T conv
197 files overlaid
CD25+ memory T conv
memory T conv
Naive T conv
Resting T reg cells
Effector Treg
count
BV785-A | ctla4
PE-Cy5.5-A | foxp3

**C**

To confirm the expression of CTLA-4 and Foxp3 in Treg cells:
Two representative samples are shown.

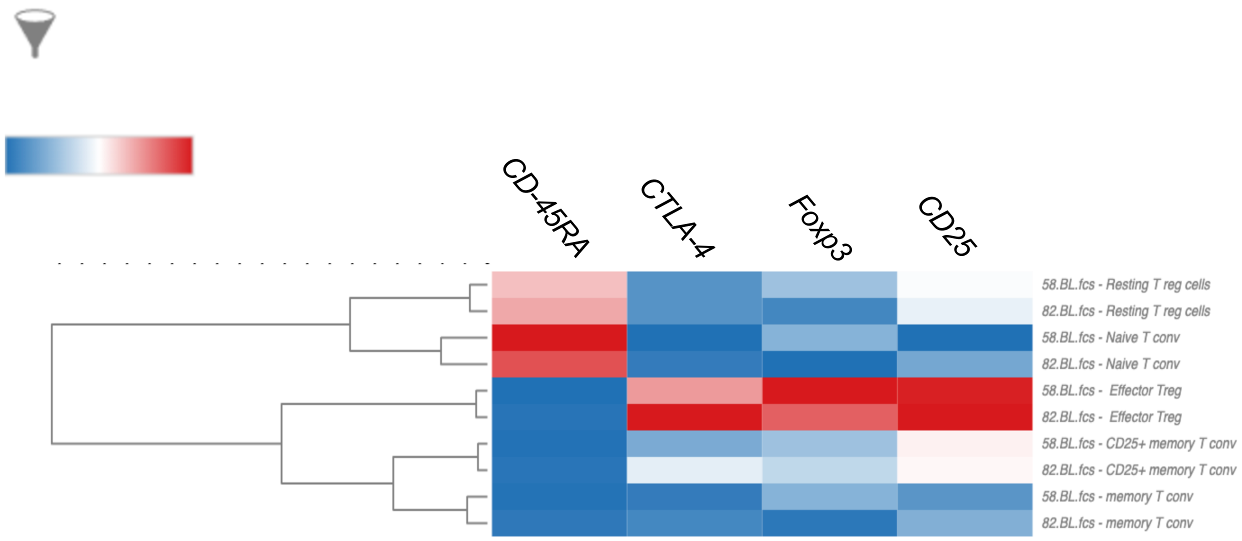


The highest expression of Foxp3 and CTLA-4 : Effector Treg cells

D

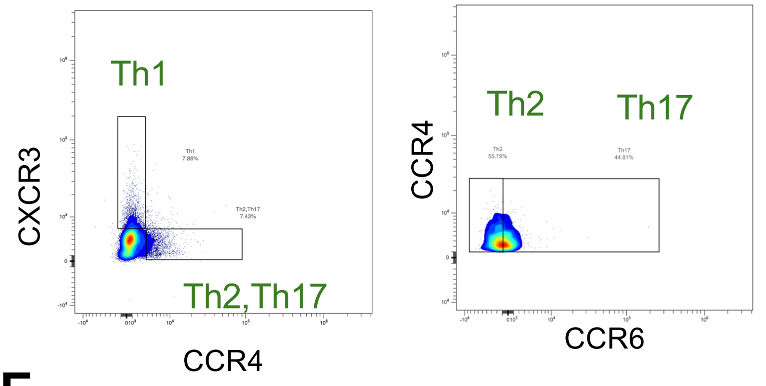
Th1
CXCR3
Th2,Th17
CCR4
Th2
Th17
CCR4
CCR6

E

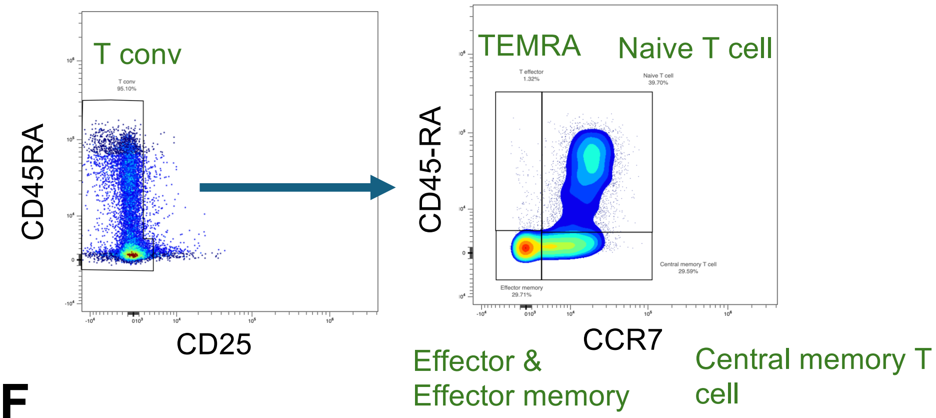
T conv
CD45RA
CD25
TEMRA
Naive T cell
CD45-RA
CCR7
Effector &
Effector memory
Central memory T
cell

F

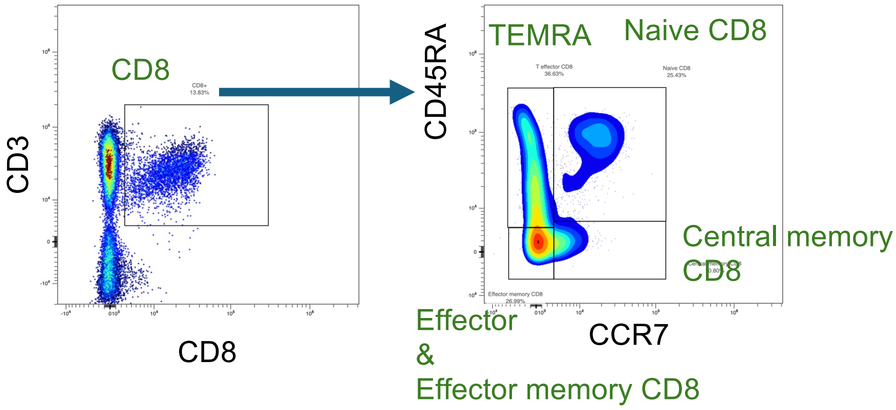
CD8
CD3
CD8
TEMRA
Naive CD8
CD45RA
Central memory
CD8
Effector
&
Effector memory CD8
CCR7

G

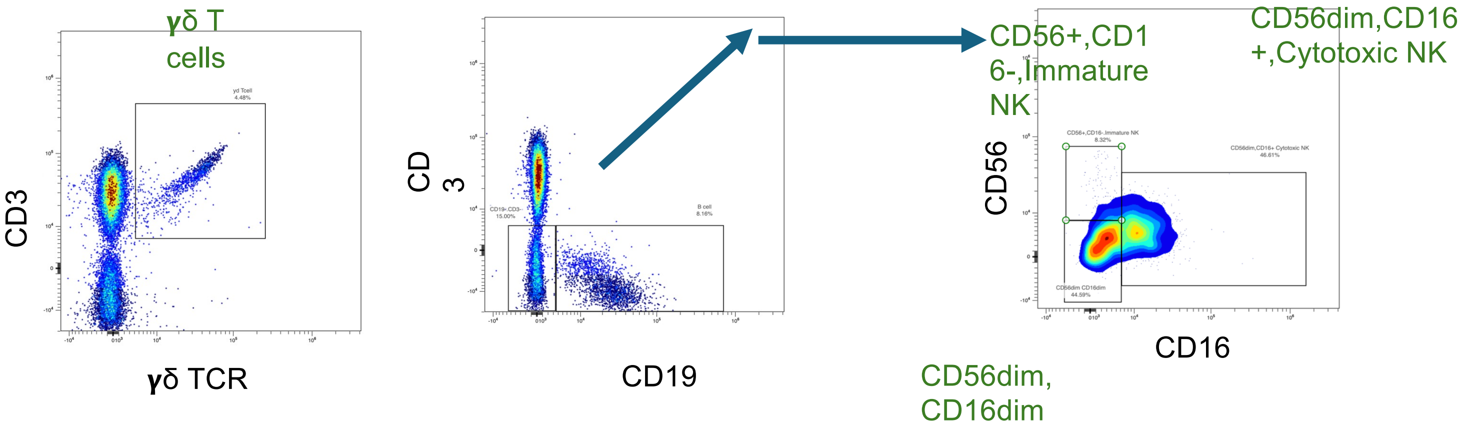
γδ T
cells
CD3
γδ TCR
CD
3
CD19
CD56+,CD1
6-,Immature
NK
CD56dim,CD16
+,Cytotoxic NK
CD56
CD16
CD56dim,
CD16dim

H

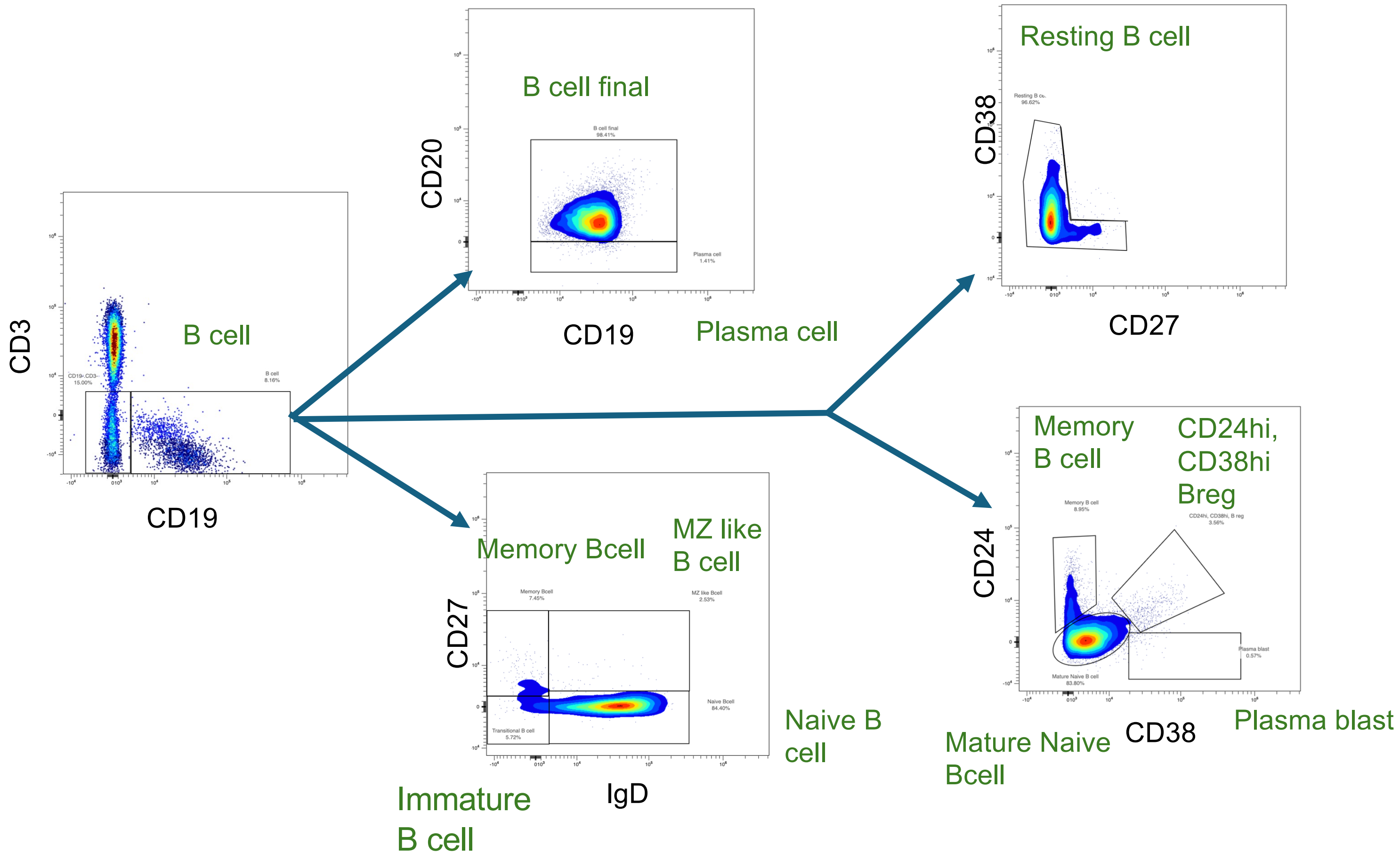

B cell
CD3
CD19
B cell final
CD20
CD19
Plasma cell
Resting B cell
CD38
CD27
Memory Bcell
MZ like
B cell
CD27
IgD
Naive B
cell
Immature
B cell
Memory
B cell
CD24hi,
CD38hi
Breg
CD24
CD38
Mature Naive
Bcell
Plasma blast

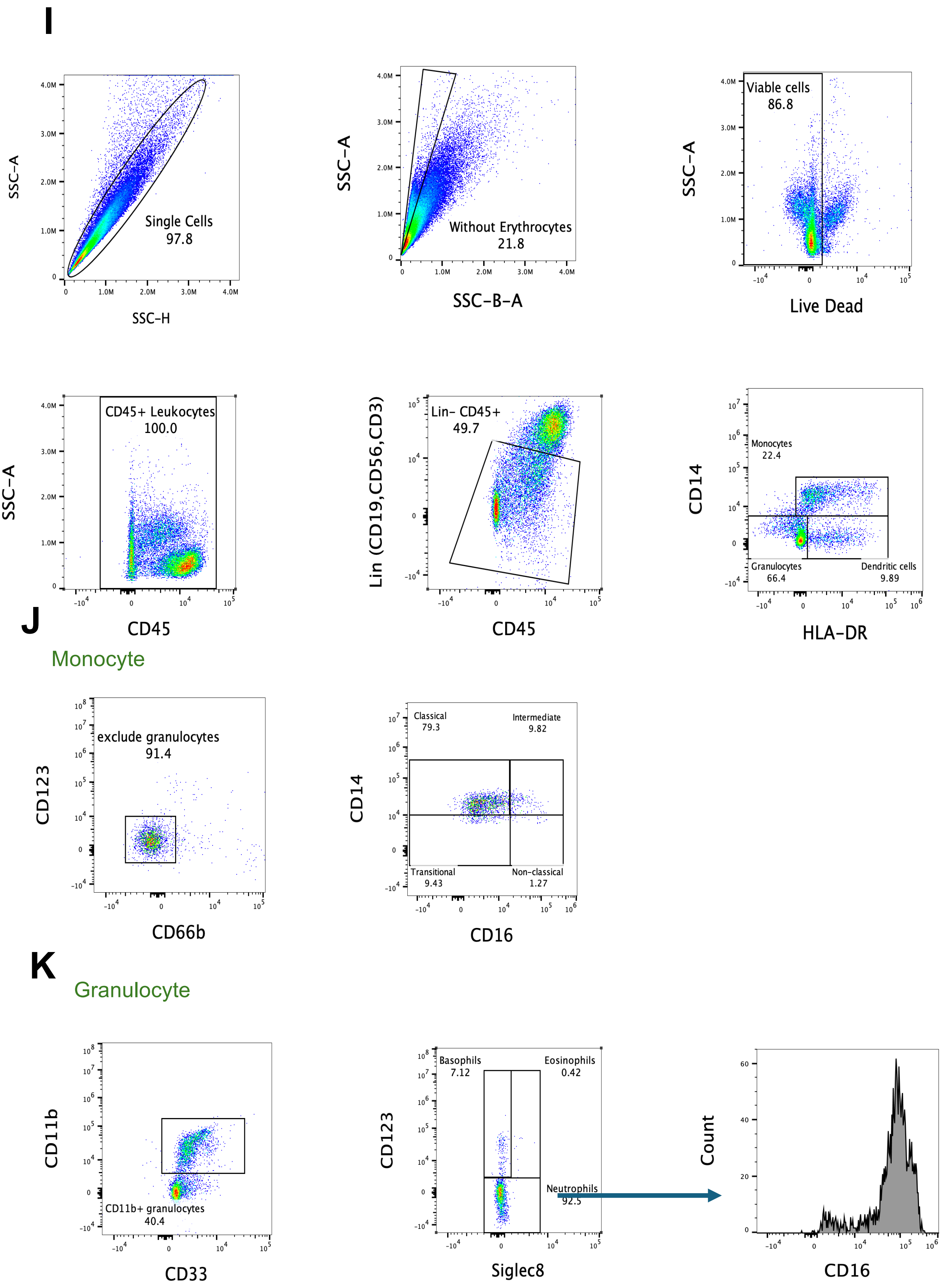
I
SSC-A
SSC-H
Single Cells
97.8
SSC-A
SSC-B-A
Without Erythrocytes
21.8
SSC-A
Live Dead
Viable cells
86.8
SSC-A
CD45
CD45+ Leukocytes
100.0
Lin (CD19,CD56,CD3)
CD45
Lin- CD45+
49.7
CD14
HLA-DR
Monocytes
22.4
Granulocytes
66.4
Dendritic cells
9.89
J
Monocyte
CD123
CD66b
exclude granulocytes
91.4
CD14
CD16
Classical
79.3
Intermediate
9.82
Transitional
9.43
Non-classical
1.27
K
Granulocyte
CD11b
CD33
CD11b+ granulocytes
40.4
CD123
Siglec8
Basophils
7.12
Eosinophils
0.42
Neutrophils
92.5
Count
CD16

**L**

#### Dendritic cell

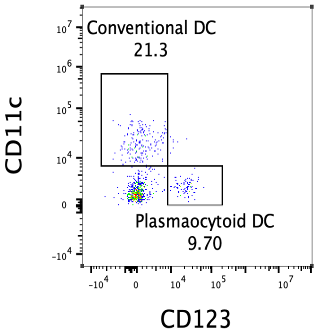


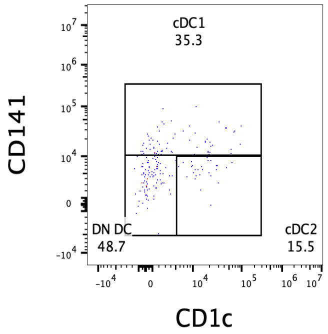


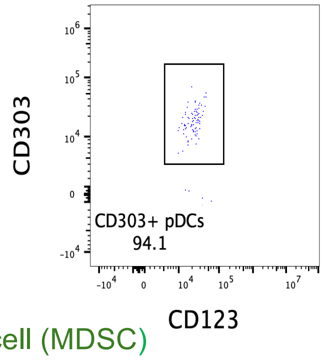


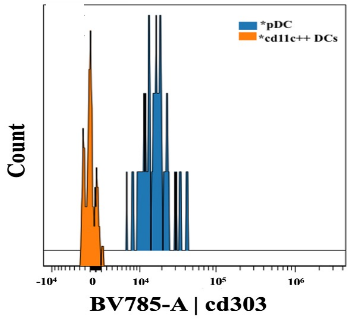


**M**

#### Myeloid-derived suppressor cell (MDSC)

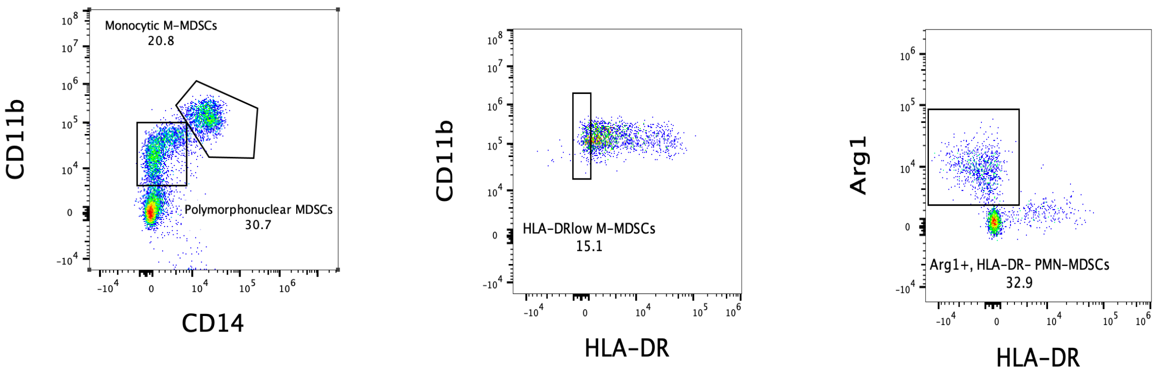

### Suppl. Fig. S5

Flowsom and unsupervised Metaclustering | Manual gating

**A**

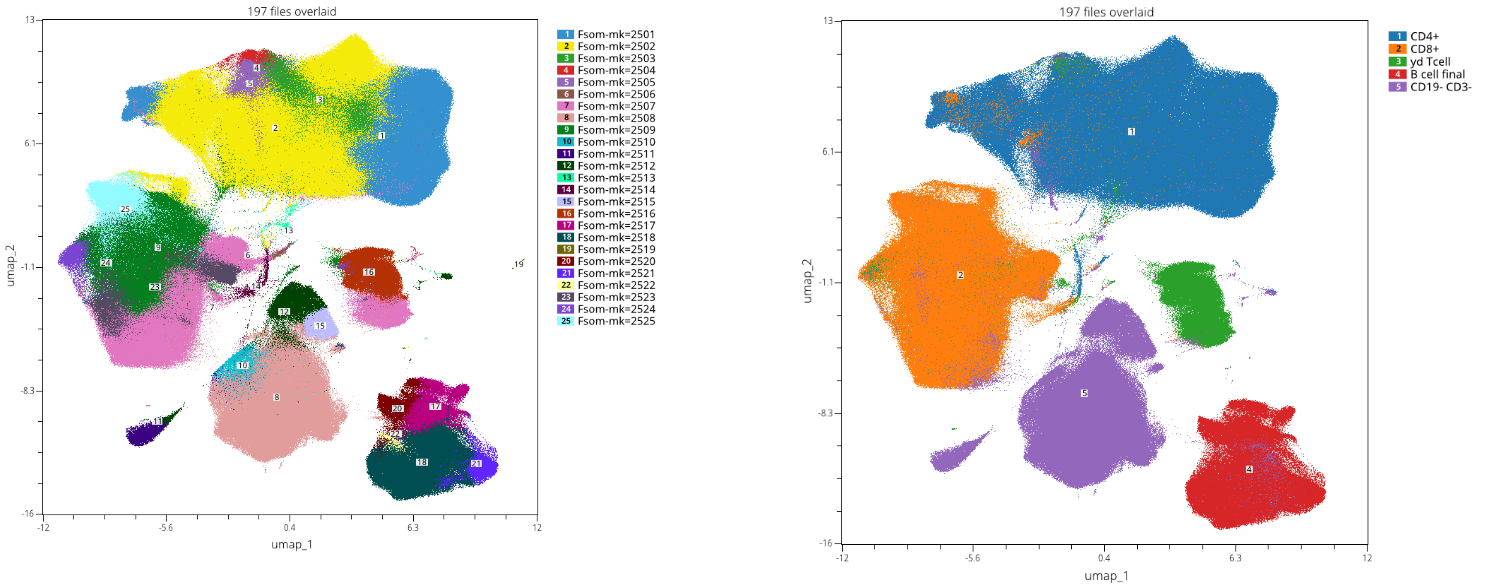


**B**

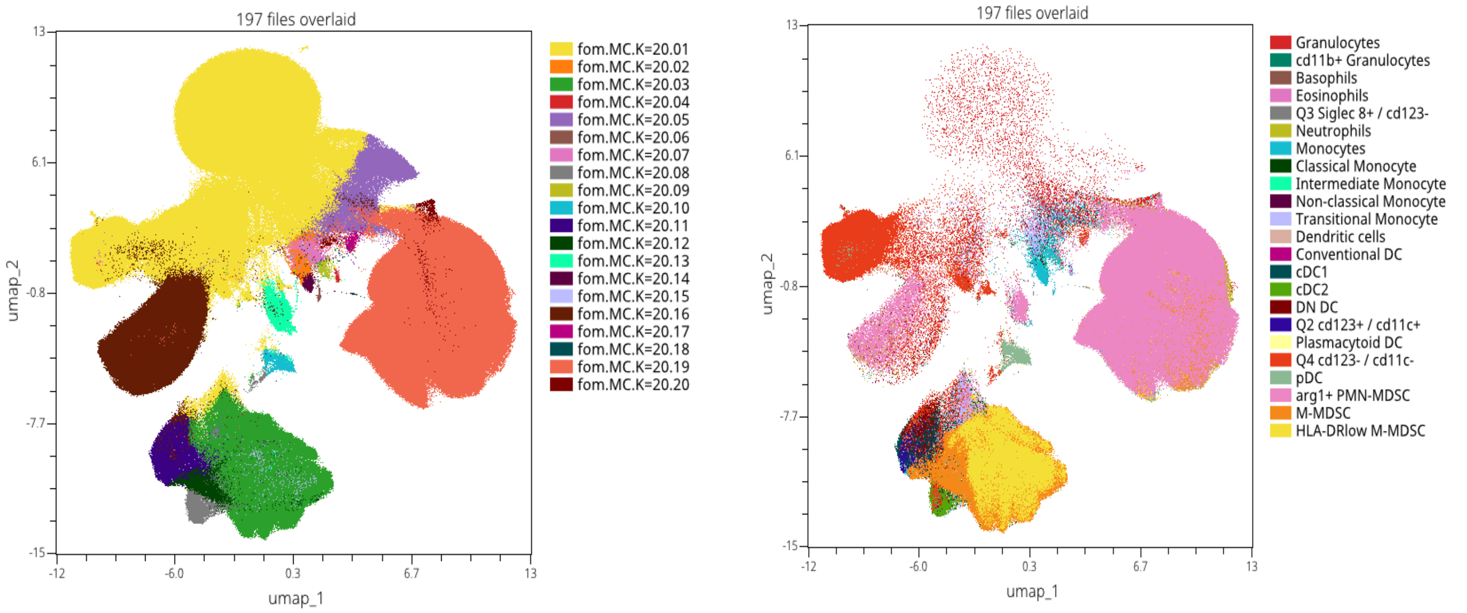

# C

#### Subsampling CD4+

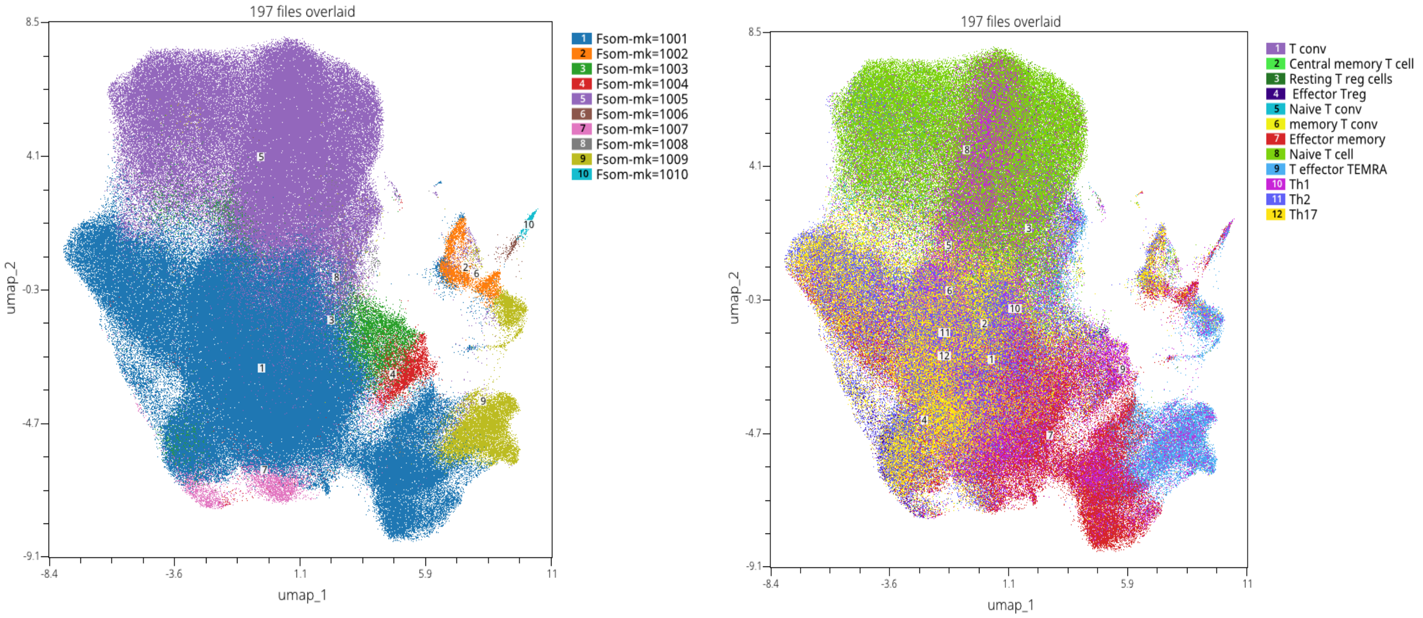


# D

#### Subsampling CD8+

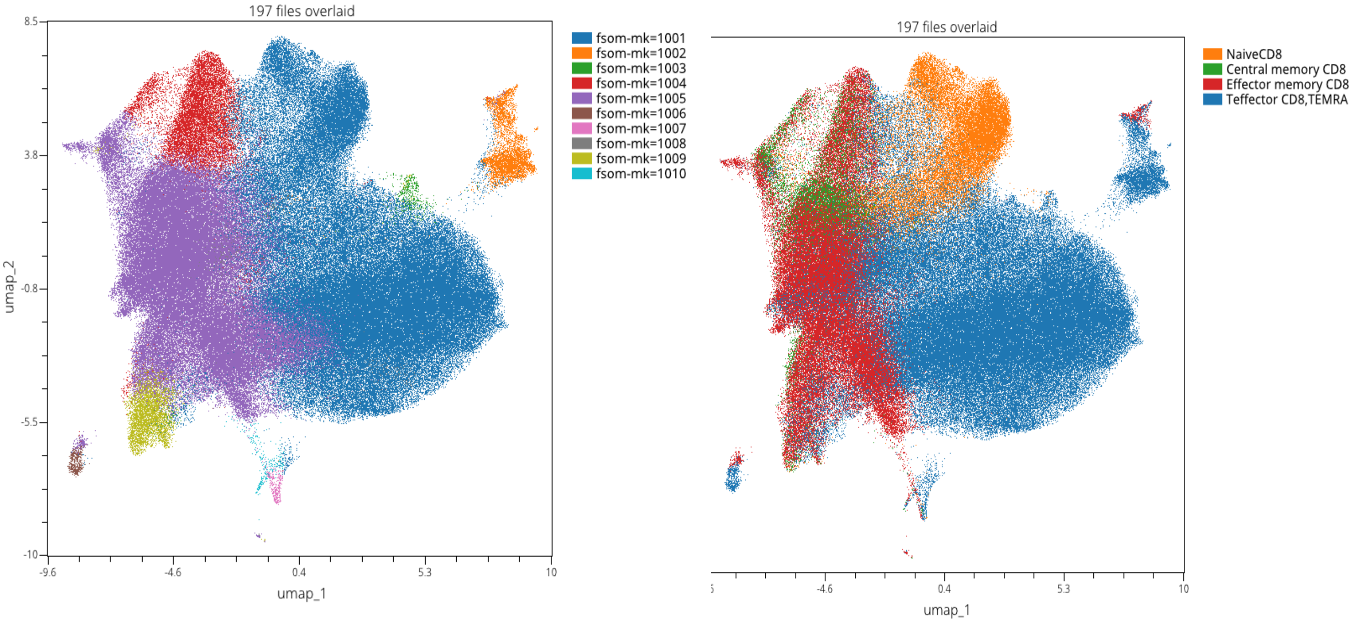

# E

#### Subsampling B cell

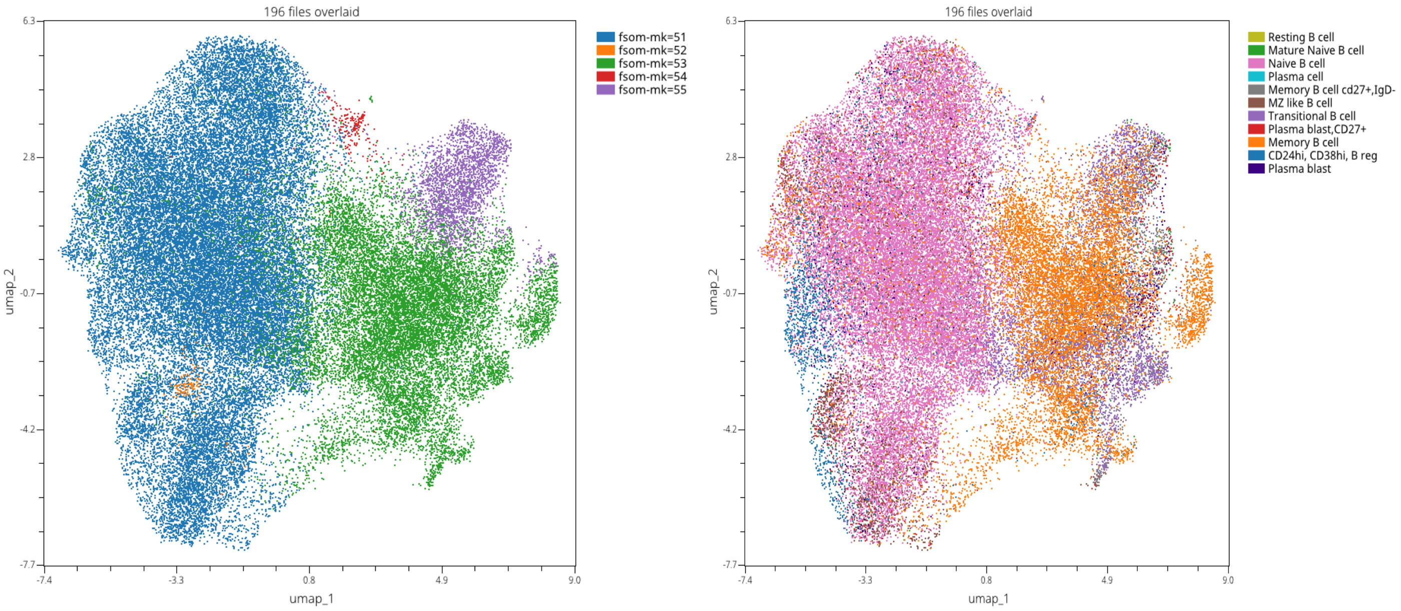


# F

#### Subsampling NK cell

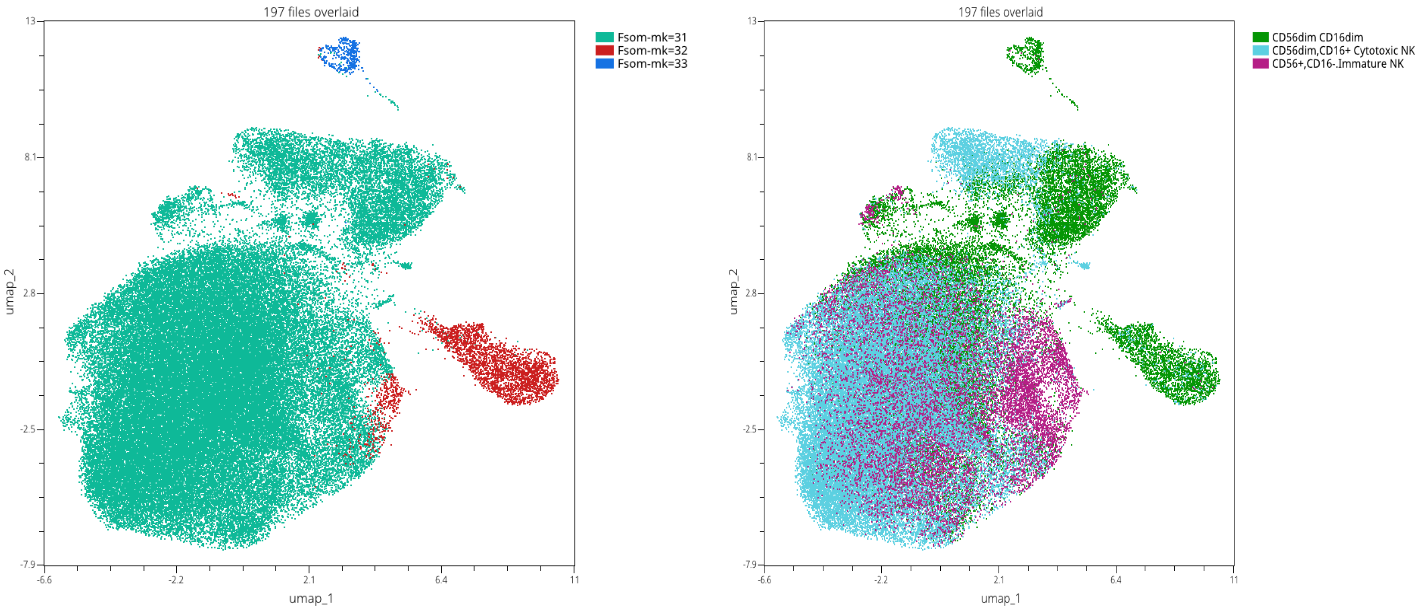

### Suppl. Fig. S6

## A

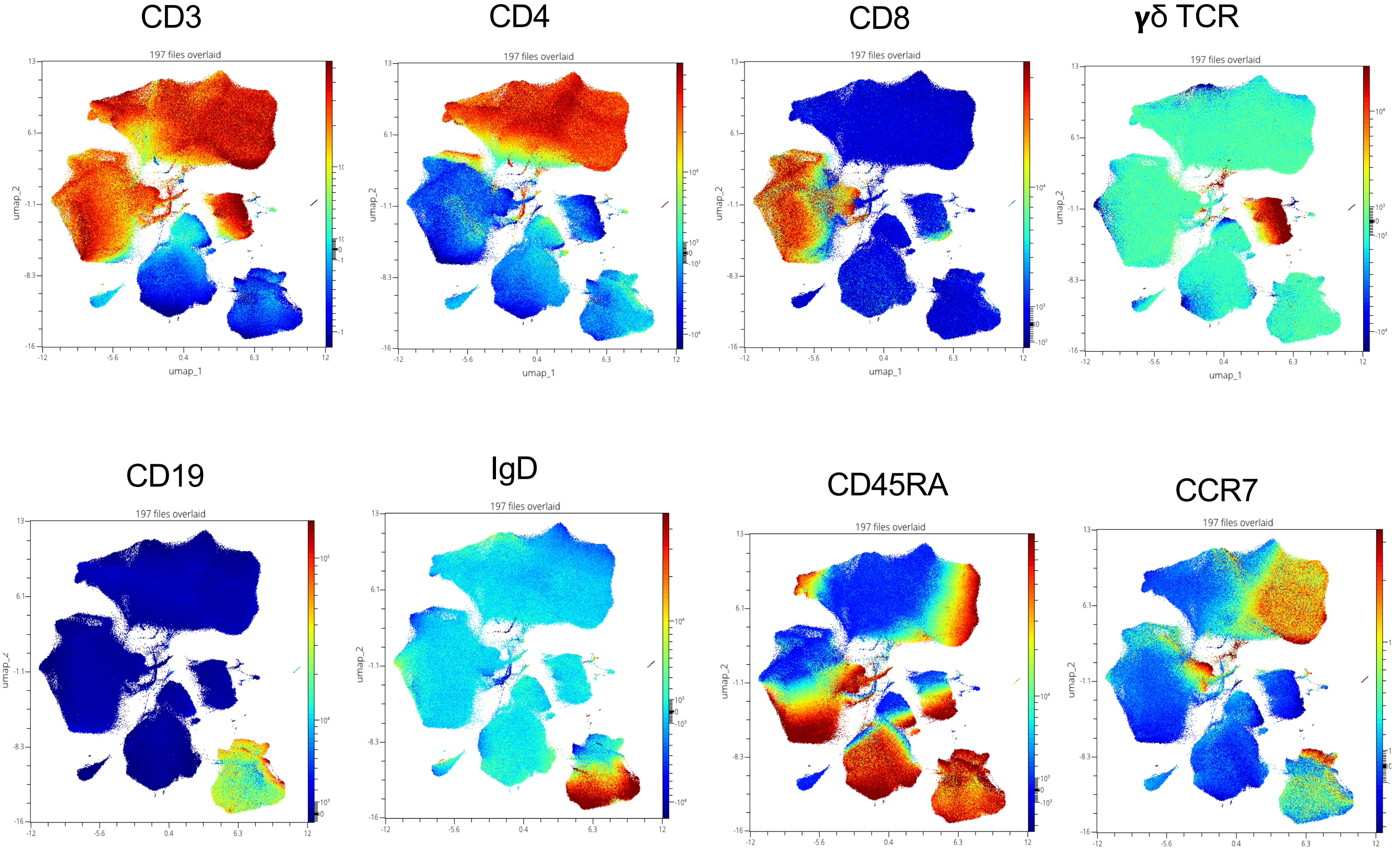


## B

Heatmap of Marker Expression across CD4⁺ Flow SOM Clusters at BL

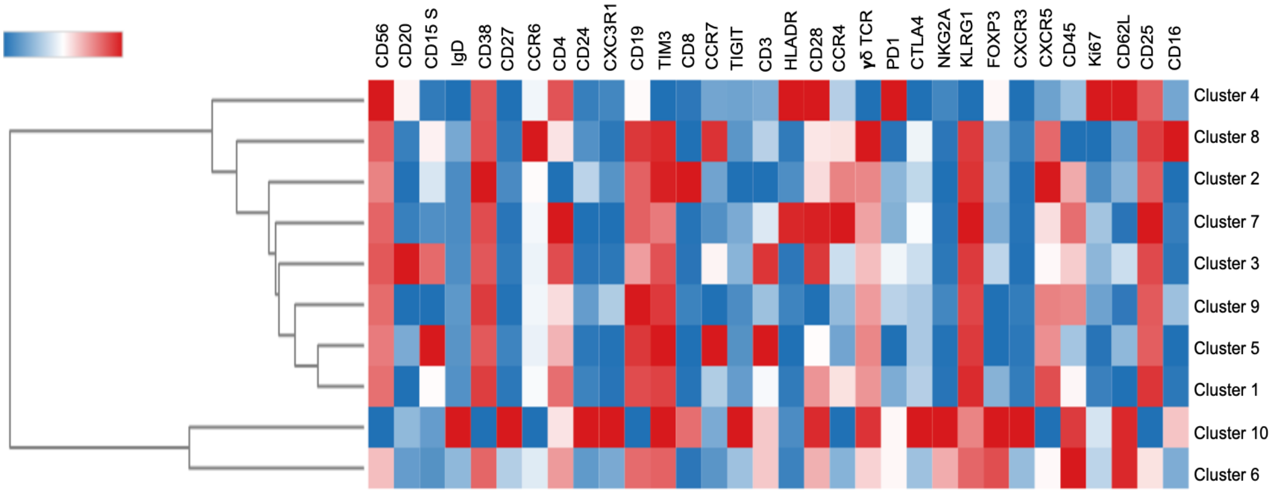

**C**

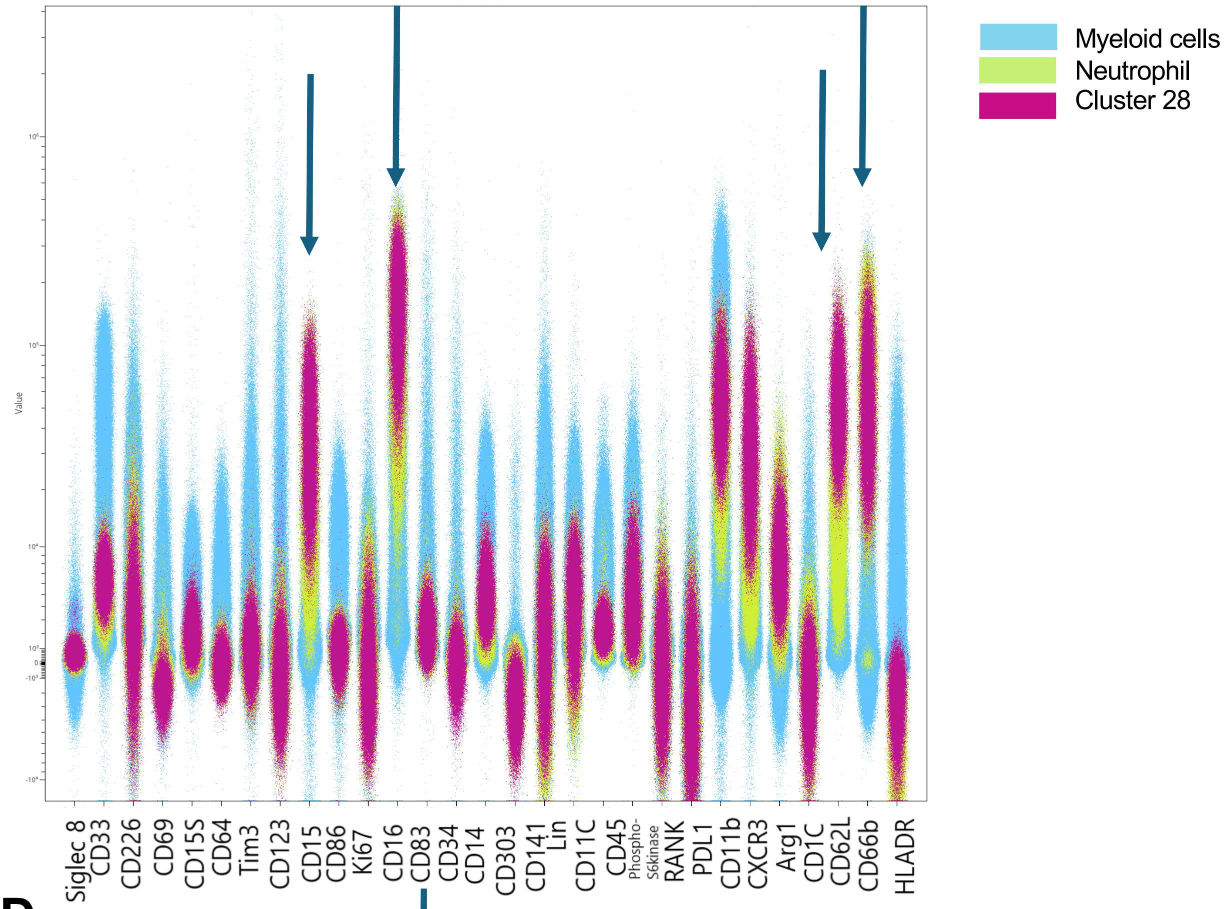
Myeloid cells
Neutrophil
Cluster 28
Value
Siglec 8
CD33
CD226
CD69
CD15S
CD64
Tim3
CD123
CD15
CD86
Ki67
CD16
CD83
CD34
CD14
CD303
CD141
Lin
CD11C
CD45
Phospho-S6kinase
RANK
PDL1
CD11b
CXCR3
Arg1
CD1C
CD62L
CD66b
HLADR

**D**

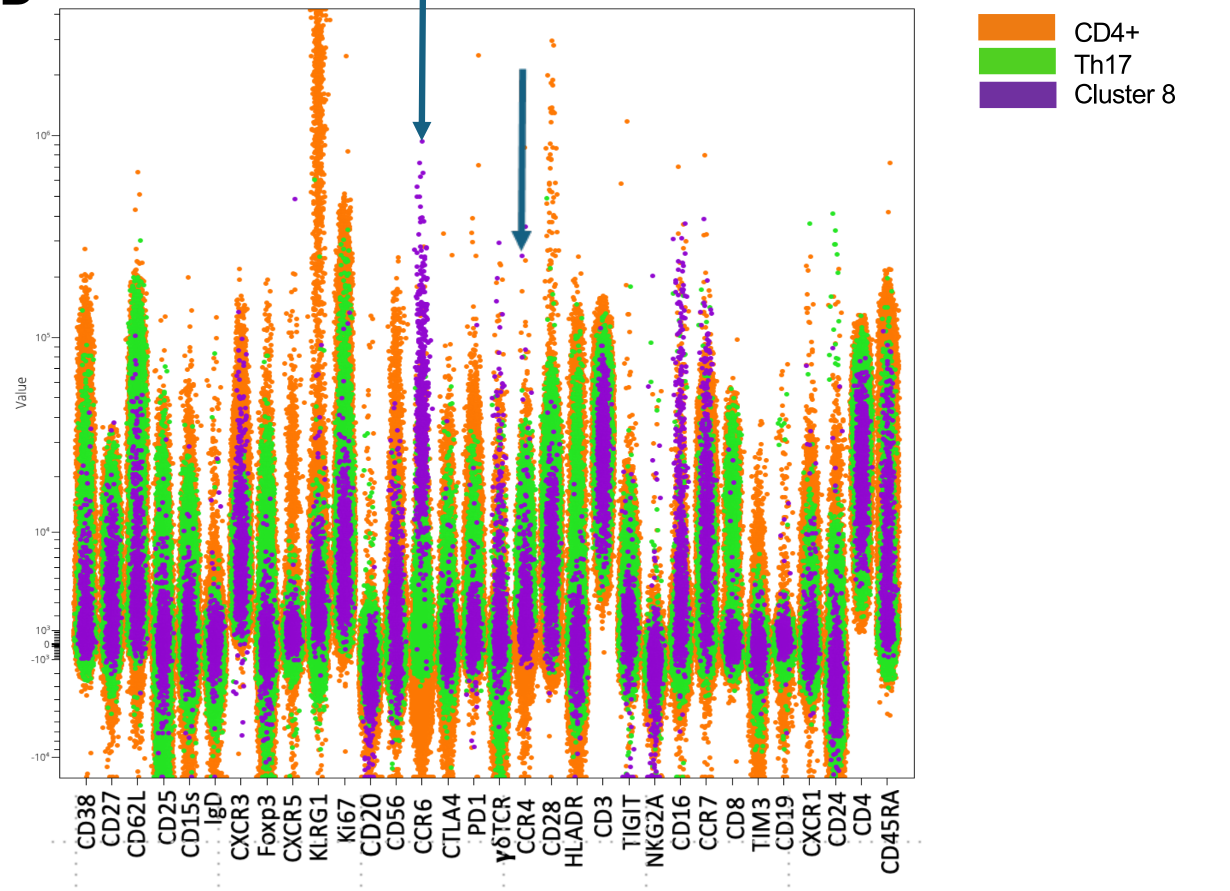
CD4+
Th17
Cluster 8
Value
CD38
CD27
CD62L
CD25
CD15s
IgD
CXCR3
Foxp3
CXCR5
KLRG1
Ki67
CD20
CD56
CCR6
CTLA4
PD1
γδTCR
CCR4
CD28
HLADR
CD3
TIGIT
NKG2A
CD16
CCR7
CD8
TIM3
CD19
CXCR1
CD24
CD4
CD45RA

### Suppl. Fig. S7

Comparative abundance of immune cell types by IFT group at baseline

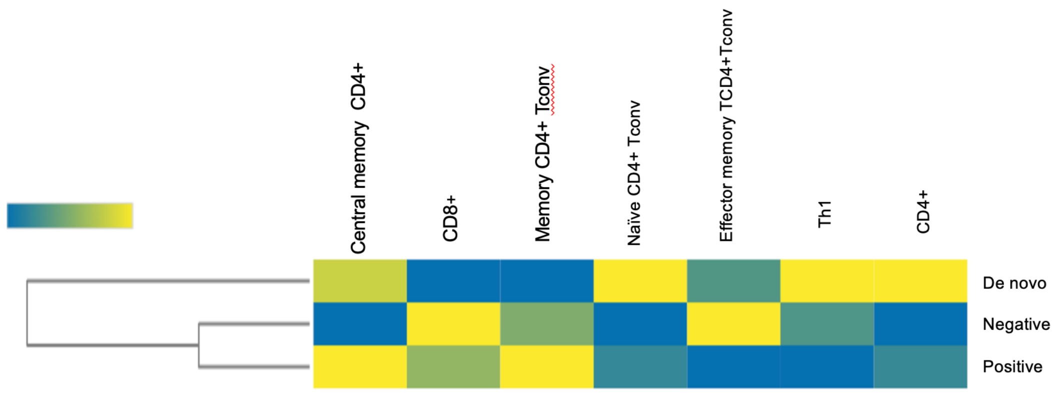

### Suppl. Fig. S8

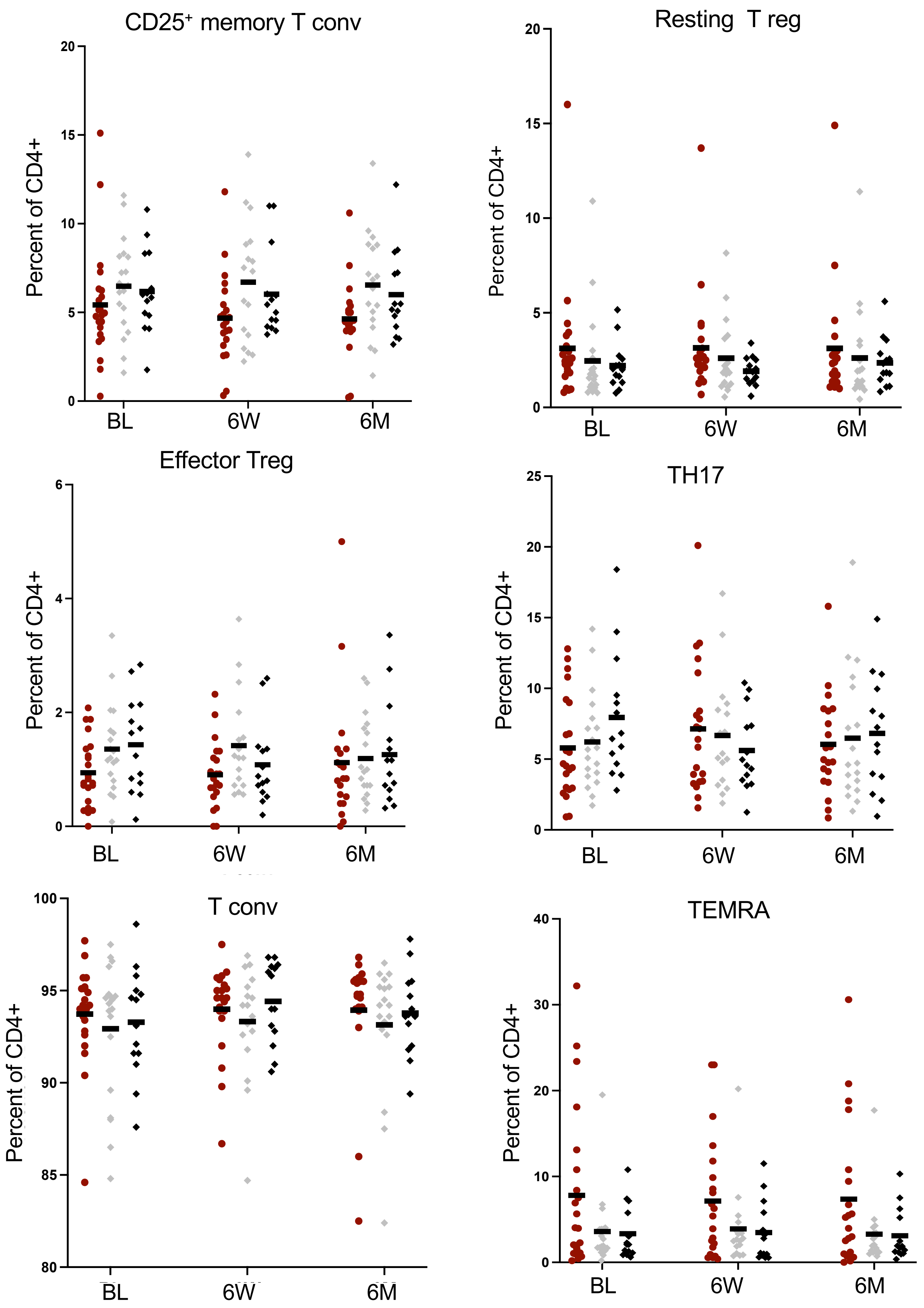
CD25⁺ memory T conv
Resting T reg
Effector Treg
TH17
T conv
TEMRA
Percent of CD4+
BL
6W
6M
0
5
10
15
20
25
2
4
6
80
85
90
95
100
30
40

### Suppl. Fig. S9

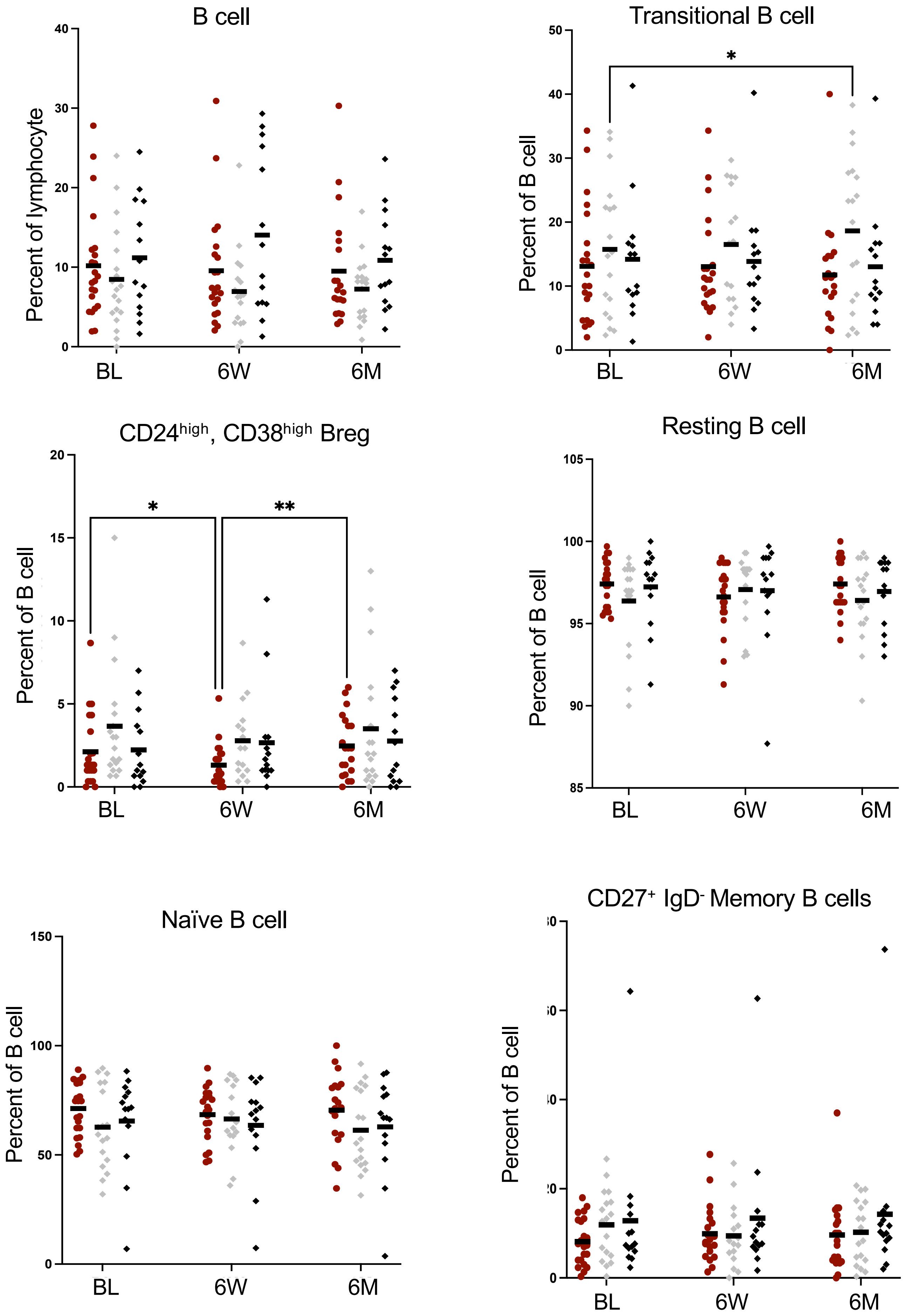
B cell
Percent of lymphocyte
0
10
20
30
40
BL
6W
6M
Transitional B cell
Percent of B cell
0
10
20
30
40
50
*
BL
6W
6M
CD24high, CD38high Breg
Percent of B cell
0
5
10
15
20
*
**
BL
6W
6M
Resting B cell
Percent of B cell
85
90
95
100
105
BL
6W
6M
Naïve B cell
Percent of B cell
0
50
100
150
BL
6W
6M
CD27+ IgD- Memory B cells
Percent of B cell
0
BL
6W
6M

Plasma blast,CD27

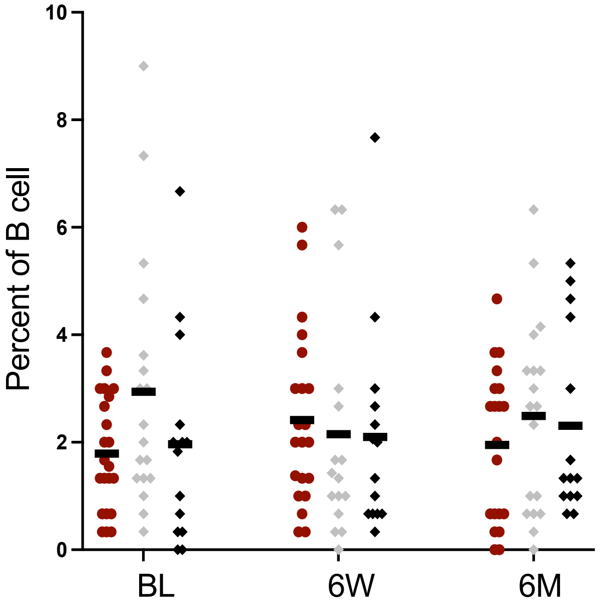


Mature Naïve B cell

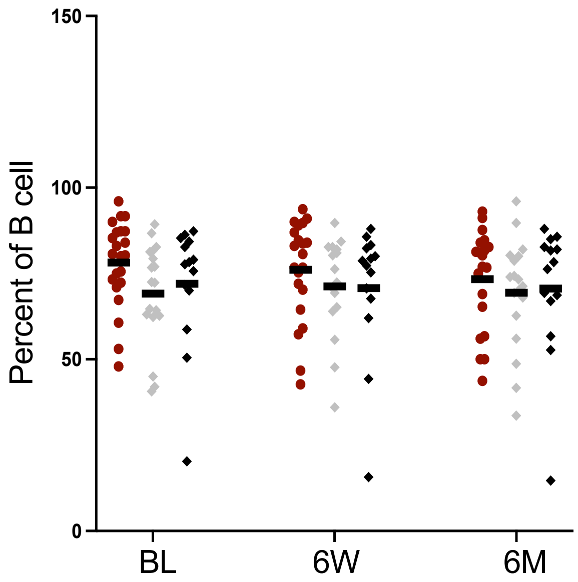


Memory B cell

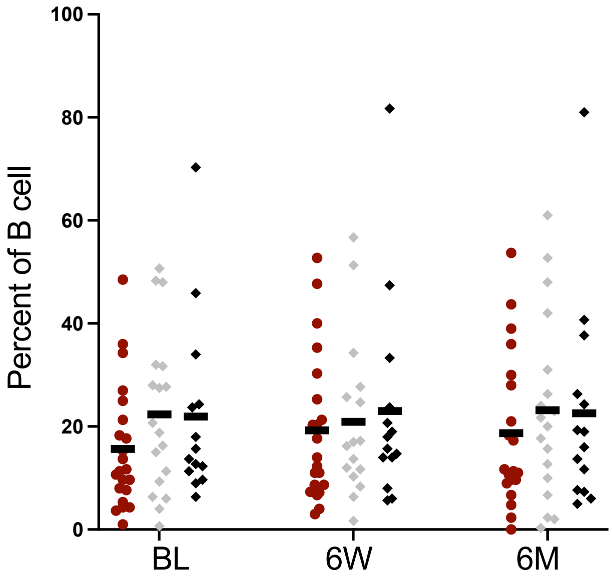


MZ like B cell

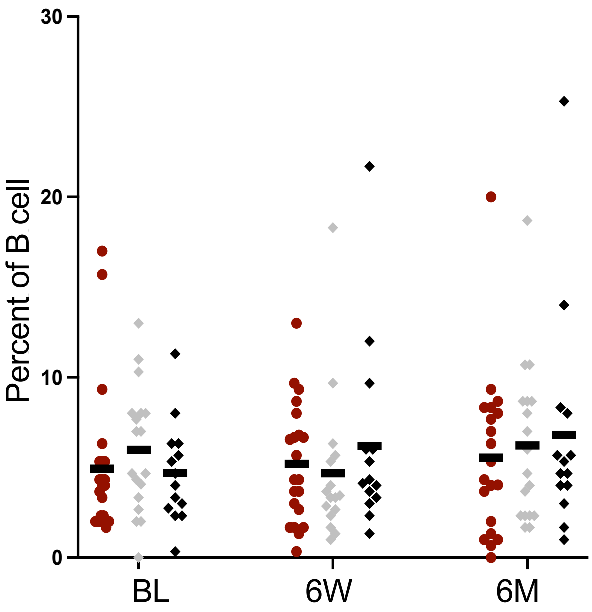

### Suppl. Fig. S10

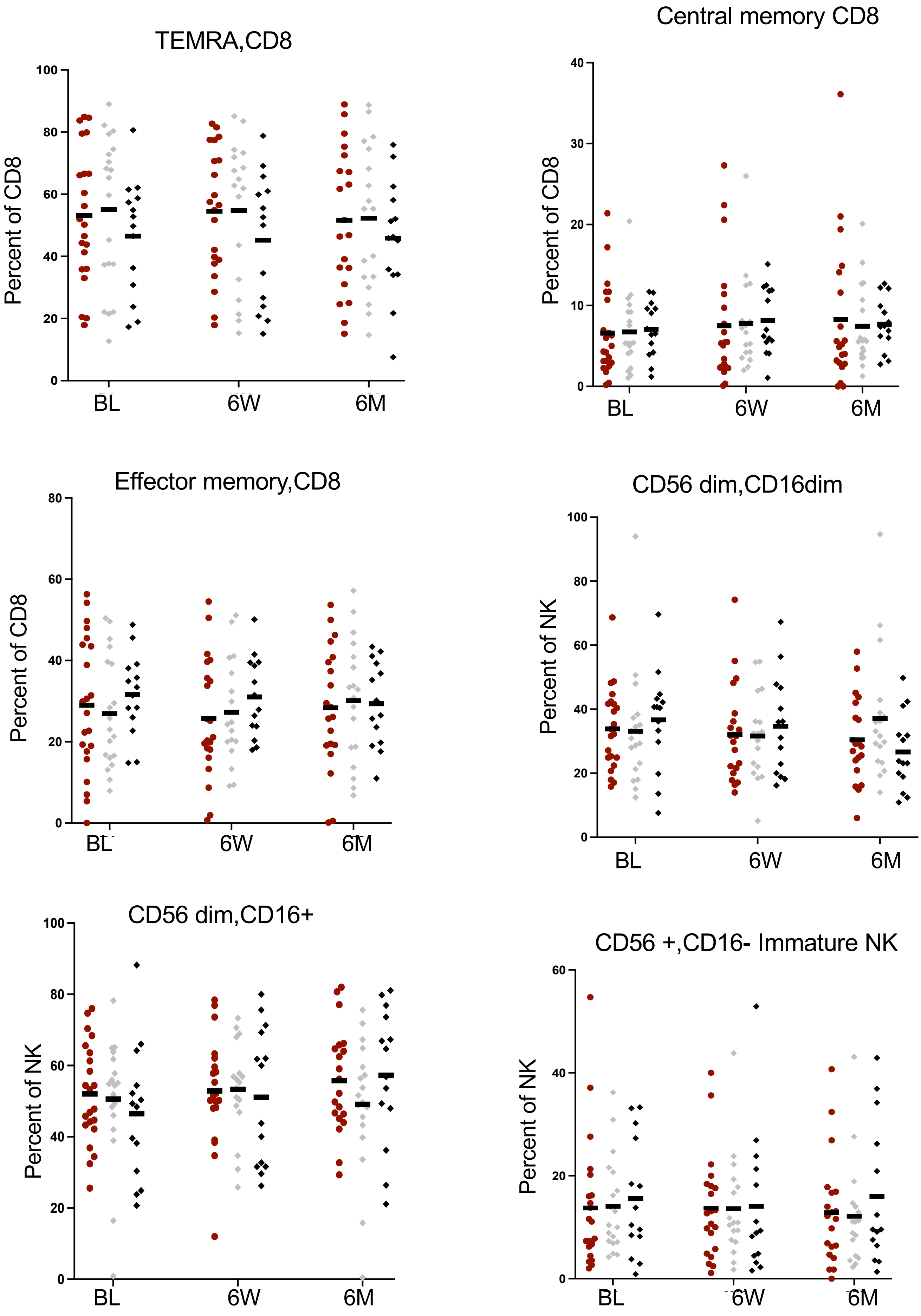

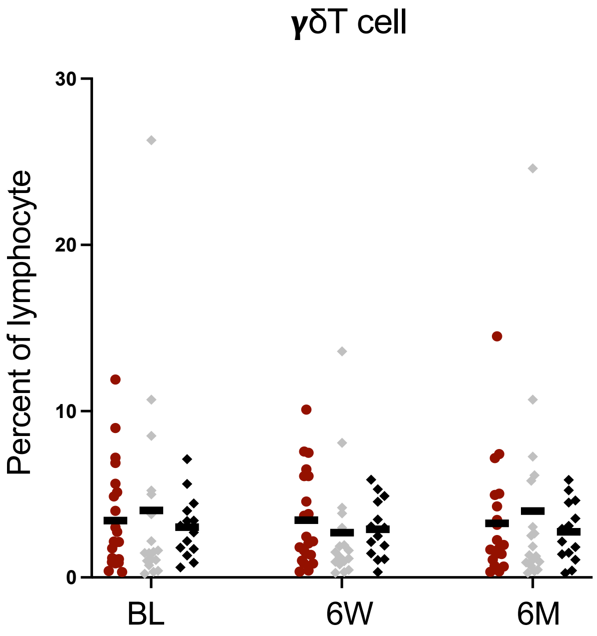
γδT cell
Percent of lymphocyte
30
20
10
0
BL
6W
6M

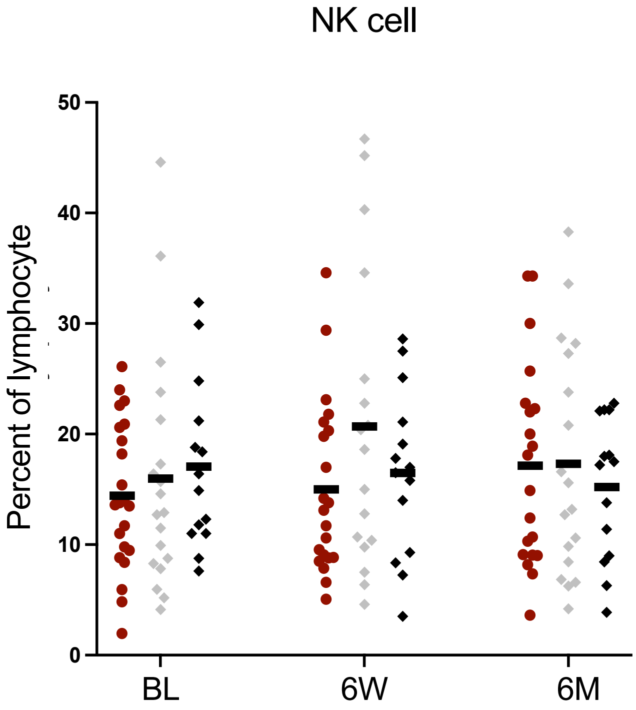
NK cell
Percent of lymphocyte
50
40
30
20
10
0
BL
6W
6M

### Suppl. Fig. 11

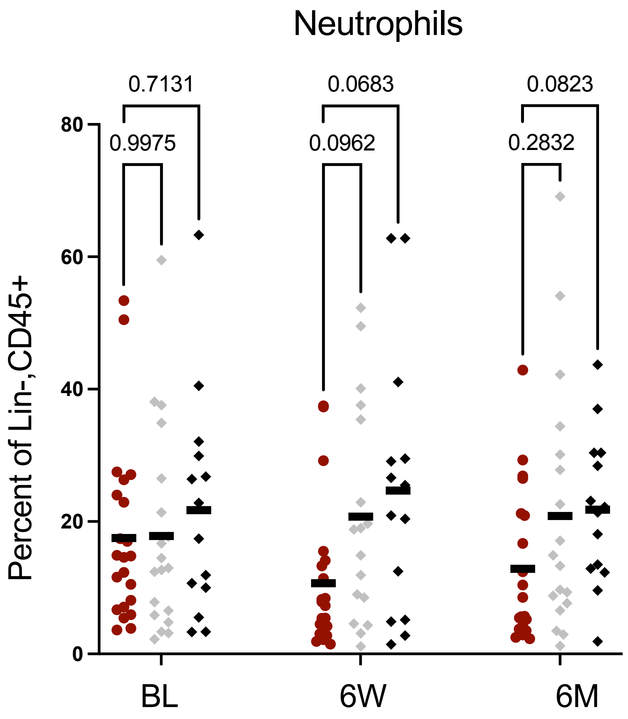
Neutrophils
0.7131
0.0683
0.0823
0.9975
0.0962
0.2832
Percent of Lin-,CD45+
80
60
40
20
0
BL
6W
6M

### Suppl. Fig. S12

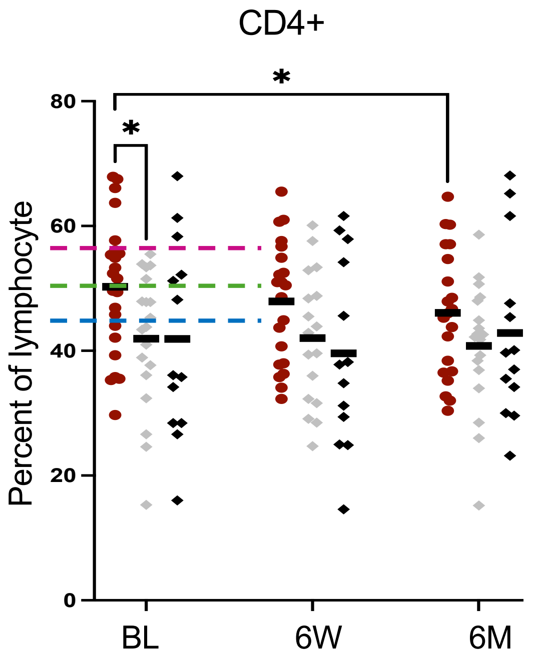


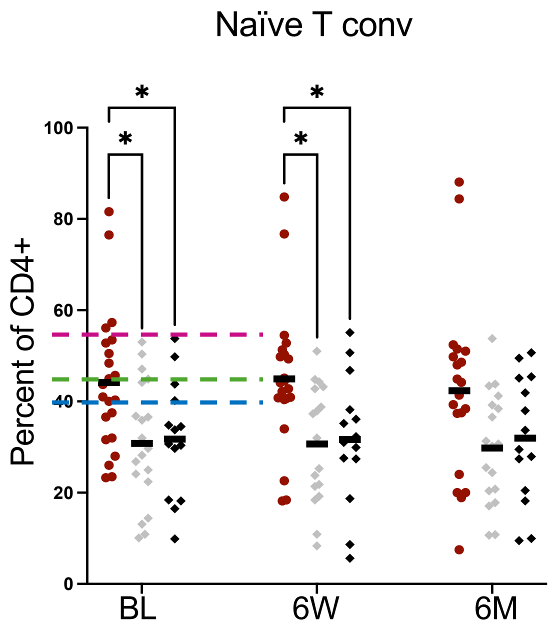


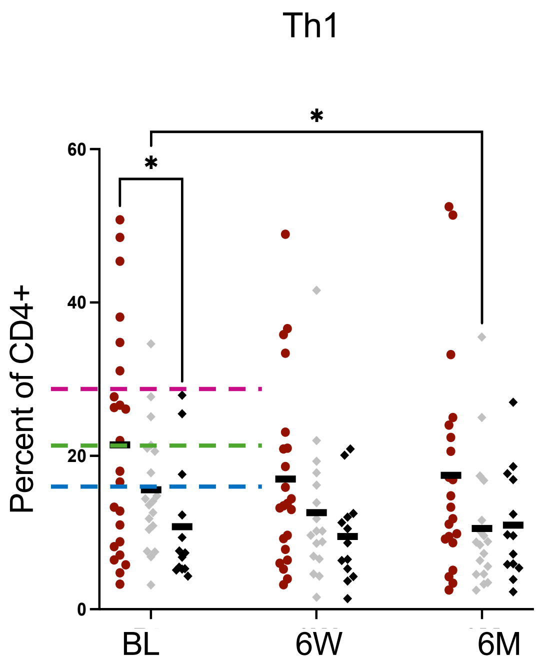


| | Likelihood ratio | *Cut off* |
|---|---|---|
| CD4+blue cut-off line | 1.7 | 45 |
| CD4+green cut-off line | 2.1 | 50 |
| CD4+pink cut-off line | Not defined | 57 |
| Naïve T conv blue cut-off line | 2.7 | 40 |
| Naïve T conv green cut-off line | 3.0 | 44 |
| Naive T conv pink cu-off line | Not defined | 55 |
| Th1blue cut-off line | 1.6 | 16 |
| Th1 green cut-off line | 3.3 | 21 |
| Th1 pink cut-off line | 5.4 | 30 |
